# Centrosome and ciliary abnormalities in fetal akinesia deformation sequence human fibroblasts

**DOI:** 10.1101/756734

**Authors:** Ramona Jühlen, Valérie Martinelli, Chiara Vinci, Jeroen Breckpot, Birthe Fahrenkrog

## Abstract

Ciliopathies are clinical disorders of the primary cilium with widely recognised phenotypic and genetic heterogeneity. Here we found impaired ciliogenesis in fibroblasts derived from individuals with fetal akinesia deformation sequence (FADS), a broad spectrum of neuromuscular disorders arising from impaired foetal movement. We show that cells derived from FADS individuals have shorter and less primary cilia (PC), in association with alterations in post-translational modifications in α-tubulin. Similarly, siRNA-mediated depletion of two known FADS proteins, the scaffold protein rapsyn and the nucleoporin NUP88, resulted in defective PC formation. Consistent with a role in ciliogenesis, rapsyn and NUP88 localised to centrosomes and PC. By proximity-ligation assays, we show that rapsyn and NUP88 are adjacent and that both proteins are adjoining to all three tubulin isoforms (α, and γ rapsyn-NUP88 interface, as well as their contact to microtubules, is perturbed in the examined FADS cells. We suggest that the perturbed rapsyn-NUP88-tubulin interface leads to defects in PC formation and that defective ciliogenesis contributes to the pleiotropic defects seen in FADS.

**Summary:** Fibroblasts derived from fetal akinesia individuals are characterised by ciliary defects and rapsyn and NUP88 are required for proper formation of the primary cilium.

## Introduction

Fetal akinesia deformations sequence (FADS), the severest form of congenital myasthenic syndromes (CMS), encompasses a broad spectrum of disorders, sharing the inability of the foetus to initiate movement (Beecroft et al., 2018; Engel, 2012; Hall, 2009; Ravenscroft et al., 2011). As a result, affected foetuses suffer from multiple joint contractures, including rocker-bottom feet, facial anomalies and lung hypoplasia. In many cases, FADS individuals are born prematurely or stillborn, and a high percentage of live-births die due to respiratory failure (Beecroft et al., 2018).

FADS and CMS can be caused by gene mutations that lead to dysfunction of the neuromuscular system, whereby in 68% of cases the defect is postsynaptic, i.e. in the skeletal muscle cells (Engel, 2012). The most frequent mutations are found in three genes, namely *RAPSN (*receptor-associated protein of the synapse, rapsyn; (Vogt et al., 2008; Winters et al., 2017)), *DOK7* (downstream of tyrosine kinase 7; (Radhakrishnan et al., 2019; Vogt et al., 2009)) and *MUSK* (muscle specific kinase; (Li et al., 2019; Tan-Sindhunata et al., 2015; Wilbe et al., 2015)), all of which are important regulators of acetylcholine receptor (AChR) formation and maintenance at the neuromuscular junction (NMJ; (Burden et al., 2018; Michalk et al., 2008)). The NMJ (also called neuromuscular synapse) is a type of synapse formed between motoneurons and the skeletal muscle fibres that, in vertebrates, use acetylcholine as neurotransmitter (Burden, 2002; Wu et al., 2010). Mutations in the subunits of the muscular nicotinic AChR have also been described in CMS and FADS (Michalk et al., 2008). We have recently expanded the spectrum of genetic causes for FADS by reporting bi-allelic, loss-of-function mutations in the nucleoporin *NUP88* as cause for a lethal form of FADS (Bonnin et al., 2018).

MuSK and rapsyn are scaffold proteins that play key roles in AChR clustering and NMJ formation. MuSK, a muscle-specific receptor tyrosine kinase, is activated by the extracellular matrix protein agrin, upon agrin-binding to Lpr4, a member of the low-density lipoprotein receptor family (Kim and Burden, 2008; Kim et al., 2008; Luiskandl et al., 2013; Mazhar and Herbst, 2012; Zhang et al., 2008; Zhang et al., 2011). Activated MuSK induces co-clustering of rapsyn and AChRs (Ghazanfari et al., 2011; Marchand et al., 2000; Marchand et al., 2002; Moransard et al., 2003), and downstream signalling of MuSK requires binding of DOK7 to the phosphotyrosine-binding site in MuSK (Okada et al., 2006). Signalling downstream of MuSK is only poorly understood, but it involves interactions of both MuSK and rapsyn to all three cytoskeleton networks, i.e. microtubules, the actin cytoskeleton and intermediate filaments (Aittaleb et al., 2015; Antolik et al., 2007; Bartoli et al., 2001; Dobbins et al., 2008; Ghasemizadeh et al., 2019; Li et al., 2016; Luiskandl et al., 2013; Mihailovska et al., 2014; Moransard et al., 2003; Oury et al., 2019). Rapsyn has, moreover, been found in non-muscle cell types (Frail et al., 1989; Musil et al., 1989) and plays a role in lysosome clustering (Aittaleb et al., 2015).

Primary cilia (PC) are discrete, non-motile microtubule-based organelles that protrude from the surface of most quiescent vertebrate cells. PC communicate extracellular signals to the cell, thereby controlling important developmental signalling pathways, such as hedgehog, Wnt, and Notch (Garcia et al., 2018; Schou et al., 2015). The core structure of the PC, the axoneme, is composed of nine, ring-forming, parallel doublets of microtubules that nucleate from the basal body and are surrounded by a ciliary membrane (Avalos et al., 2017). Doublets consist of a complete A-tubule scaffolding a partial B-tubule (Mirvis et al., 2018). Within the PC, tubulin is subjected to distinct post-translational modifications, in particular acetylation, glutamylation/detyrosination, and glycylation, all of which control PC assembly and length (Gadadhar et al., 2017; Gaertig and Wloga, 2008; Keeling et al., 2016). Additionally, the PC requires a bidirectional transport system, known as intraflagellar transport (IFT), that transfers and correctly localises proteins required for PC growth, maintenance and signalling (Avalos et al., 2017). The A-tubule binds the IFT retrograde motor protein dynein, and the B-tubule binds the IFT anterograde motor protein kinesin-II (Mirvis et al., 2018). Besides IFT, two additional, IFT-independent routes within the lumen of PC are known: passive diffusion and vesicle trafficking (Luo et al., 2017; Ruba et al., 2019). The ciliary proteome is composed of more than 1300 proteins and about 52 subcomplexes (Boldt et al., 2016). Defects in function or structure of the PC give rise to pleiotropic genetic disorders and manifestations include brain malformations, facial anomalies, neurodevelopmental disorders, such as Joubert syndrome, congenital heart defects, trisomy 21, and skeletal malformations (Galati et al., 2018; Reiter and Leroux, 2017; Wang and Dynlacht, 2018).

Here, we provide evidence that defects in ciliogenesis contribute to the pleiotropic defects seen in FADS.

## Results

### Nuclei of FADS fibroblasts are frequently misshapen and lobulated

While genetic causes and clinical features of FADS are relatively well described, cellular consequences of pathogenic variants in FADS-related genes are largely unknown. We therefore pursued a cell biological characterisation of fibroblasts derived from two FADS individuals: one with unknown genetic cause (FADS 1) and a second one with a homozygous c.484G>A (p.Glu162Lys) variant of *RAPSN* (FADS 2; *(Winters et al., 2017)*). Confocal microscopy revealed that fibroblasts derived from these individuals, in contrast to normal human foetal fibroblasts (MRC5), often had nuclei with abnormal shape and a lobulated nuclear envelope (NE), similar to fibroblasts from Hutchinson-Gilford progeria syndrome (HGPS) patients (Fig. 1A, B). About 25-30% of FADS cells exhibited deformed nuclei compared to 50% in case of HGPS, and only 10% of MRC5 cells (Fig. 1C). Nuclear shape and contour irregularities were visualised by immunofluorescence studies with antibodies against lamin A/C (LA/C; Fig. 1A) and lamin B1 (LB1; Fig. 1B). Note the intranuclear lamin B1 puncta in all three fibroblast lines as previously described in other cell lines (Drozdz et al., 2017; Jevtic et al., 2019; Stephens et al., 2018). Despite these similar variations in nuclear morphology in FADS and HGPS fibroblasts, expression of lamin A/C and lamin B1 in FADS fibroblasts was only slightly altered, whereas all three lamin isoforms were significantly reduced in HGPS cells in comparison to MRC5 cells (Fig. 1D, E).

**Figure 1:**
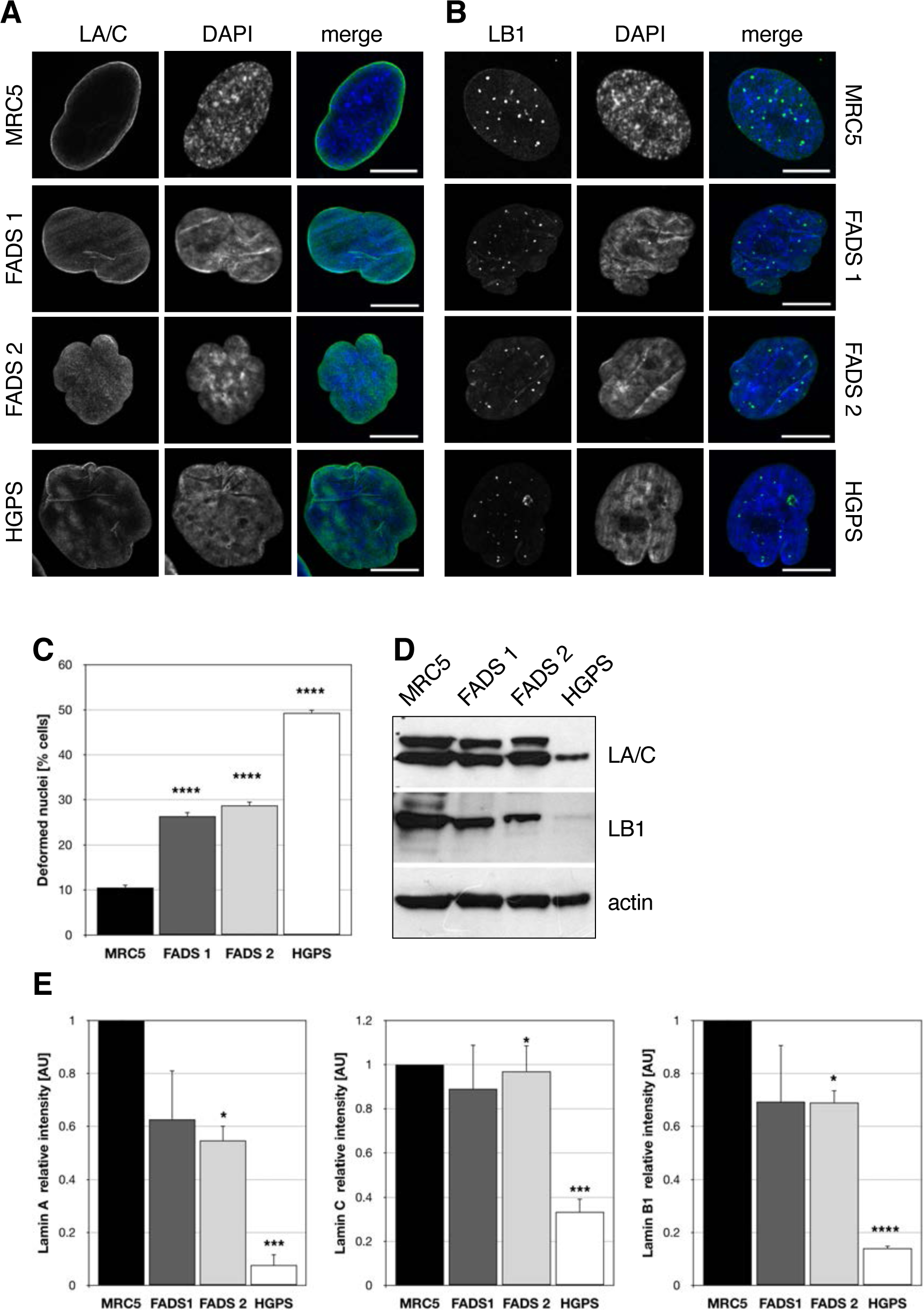
Cultured fibroblasts from FADS individuals show changes in nuclear morphology. Representative confocal images of cultured fibroblasts from a healthy control foetus (MRC5), two FADS individuals, and a Hutchison Gilford progeria syndrome (HGPS) patient were stained for (**A**) lamin A/C (LA/C, green) and (**B**) lamin B1 (LB1, green) and percentage of deformed nuclei was quantified based on LB1 staining (**C**). DNA was visualised with DAPI (blue). Scale bars, 5 µm. (**D**) Western blot analysis of LA/C and LB1 expression levels from cultured control and FADS fibroblasts and their densitometric quantification (**E**). Actin was used as loading control. P-values ****<0.0001, ***<0.001, *<0.05, t-test, two-tailed.

### FADS fibroblasts are less proliferative

To further characterise FADS fibroblasts in comparison to normal foetal fibroblasts, we subsequently analysed the proliferation of these cells. First, we determined the proliferation rate of cultured FADS fibroblasts in comparison to MRC5 fibroblasts. As shown in Fig. 2A, growth rates of FADS fibroblasts were significantly decreased compared to the MRC5 control cells: the growth constant k, i.e. slope of the linear regression line, was 0.0039 for FADS 1, 0.0050 for FADS 2, and 0.0079 for MRC5. Next, we defined the percentage of cells positive for the proliferation marker Ki-67. Ki-67 marks cells in all active phases of the cell cycle, i.e. G1, S, G2, and M phase (Gerdes et al., 1984). As shown in Fig. 2B, about 55% of MRC5 and ∼60% of FADS 2 fibroblasts were in the active phases of the cell cycle, while less than 40% of FADS 1 cells were Ki-67 positive. To more specifically determine the proliferation rate, we analysed DNA synthesis by incorporation of 5-ethynyl-20-deoxyuridine (EdU), a thymidine analog which is incorporated into newly synthesised DNA upon replication (Buck et al., 2008). EdU incorporation revealed that DNA synthesis in both FADS fibroblast lines was significantly reduced as compared to MRC5 cells (Fig. 2C). The reduced proliferation rate did only coincide with a marked increase in cellular senescence in FADS 1 fibroblasts, as revealed by β-galactosidase staining (Fig. 2D). Flow cytometry further revealed that the overall cell cycle distribution was similar in FADS and control fibroblasts (Fig. 2E). Together these data indicate that FADS fibroblasts have a reduced proliferation rate, but not necessarily a reduced proliferative state.

**Figure 2:**
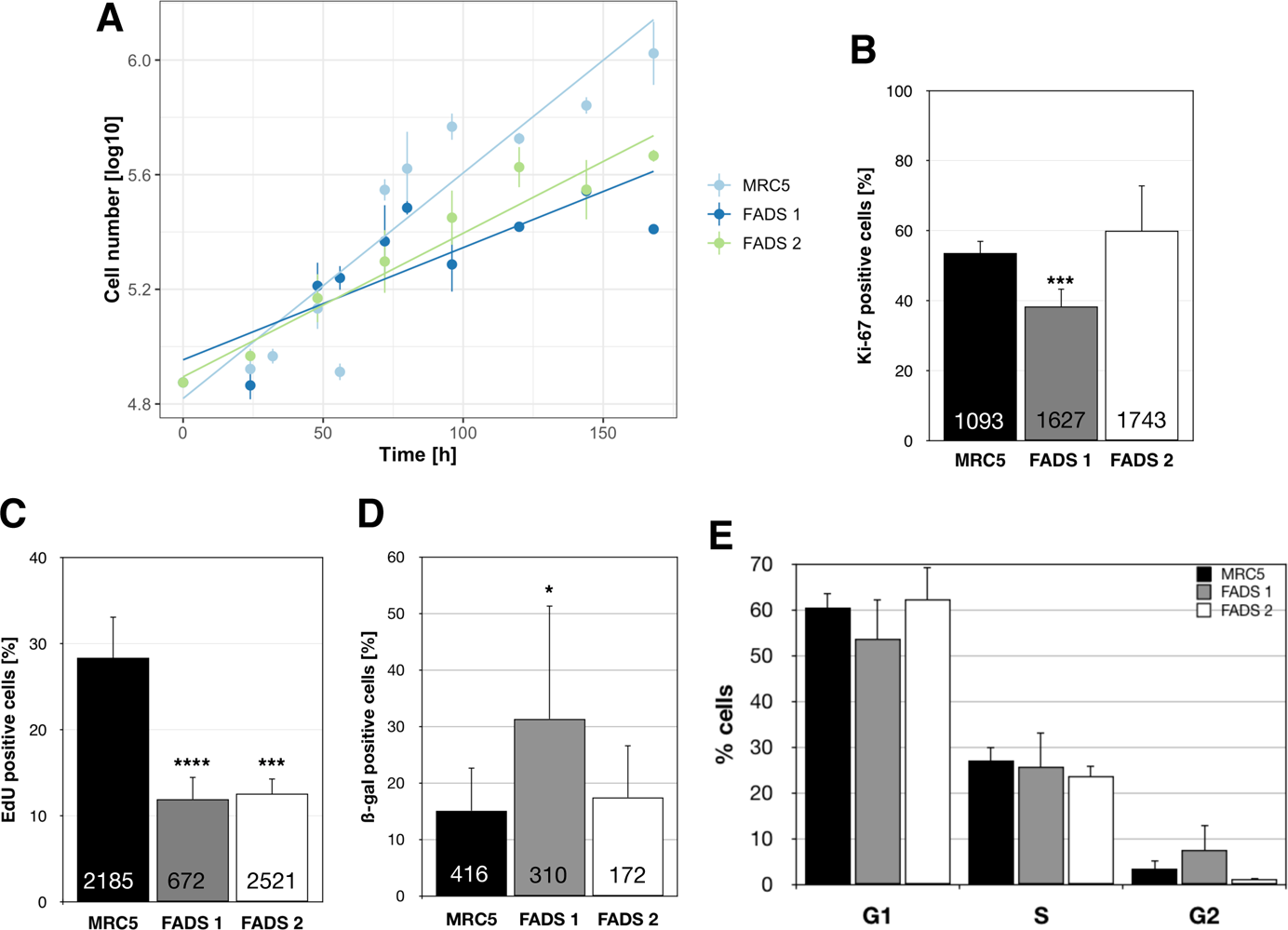
FADS fibroblasts exhibit a decreased proliferative state. (**A**) Equal cell numbers were seeded 24 h before measurement and cell numbers were determined at the indicated time points. Growth curve analysis and growth constant k (slope of regression curve) calculation was done using multilevel regression technique using R. Data points represent the mean of at least three independent experiments at the indicated time point and the point range shows the SEM. Growth constants k (slope of linear regression line) are decreased in both FADS cell lines compared to control MRC5 cells. P-values < 0.001, Pearson’s χ test. (**B**) The proliferative state of the indicated fibroblast cell line was determined by immunofluorescence microscopy and anti-Ki-67 staining. The proliferative state of FADS 1, but not FADS 2 cells, is reduced as compared to MRC5 control cells. (**C)** 5-ethynyl-20-deoxyuridine (EdU) incorporation and microscopic analysis revealed that the respective proliferation rate of both FADS 1 and FADS 2 cells was significantly lower than of MRC5 cells. (**D**) Cellular senescence was evaluated by β-galactosidase staining in FADS and MRC5 fibroblasts. (**E**) Cell cycle distribution of MRC5 and FADS fibroblasts revealed by flow cytometry. Data present the mean ±SD of at least three independent experiments. Total number of analysed cells is indicated at the bottom of each bar. P-values ****<0.0001, ***<0.001, *<0.05, t-test, two-tailed.

### Centrosomal and microtubule nucleation anomalies in FADS fibroblasts

In order to better understand proliferative defects that coincide with FADS and given the link of rapsyn (and MuSK) to the cytoskeleton, we next examined properties of microtubules (MTs) and mitotic spindles in FADS cells. After cold-treatment, we observed abnormal sloppy tubulin spindles in late stages of mitotic FADS 1 fibroblasts (Fig. S1A), whereas the overall stability of MTs using high and low concentrations of nocodazole was not altered in FADS 1 fibroblasts (Fig. S1B). When monitoring the regrowth of MTs after cold-induced depolymerisation, regrowth from the centrosome near the NE was observed within 30 s of rewarming to 37°C in MRC5 cells (Fig. 3A). In FADS 1 and FADS 2 cells, however, MT regrowth was initiated at random sites in the cytoplasm during the first 30 s of rewarming. After 90 s both FADS 1 and FADS 2 fibroblasts showed increased MT polymerisation/growth speed (Fig. 3A, 90 s). Non-centrosomal (nc) MT polymerisation was observed in less than 10% of MRC5 cells, whereas 70-80% of FADS 1 and FADS 2 showed both, centrosomal and ncMT regrowth (Fig. 3B, 30s and 90s).

**Figure 3:**
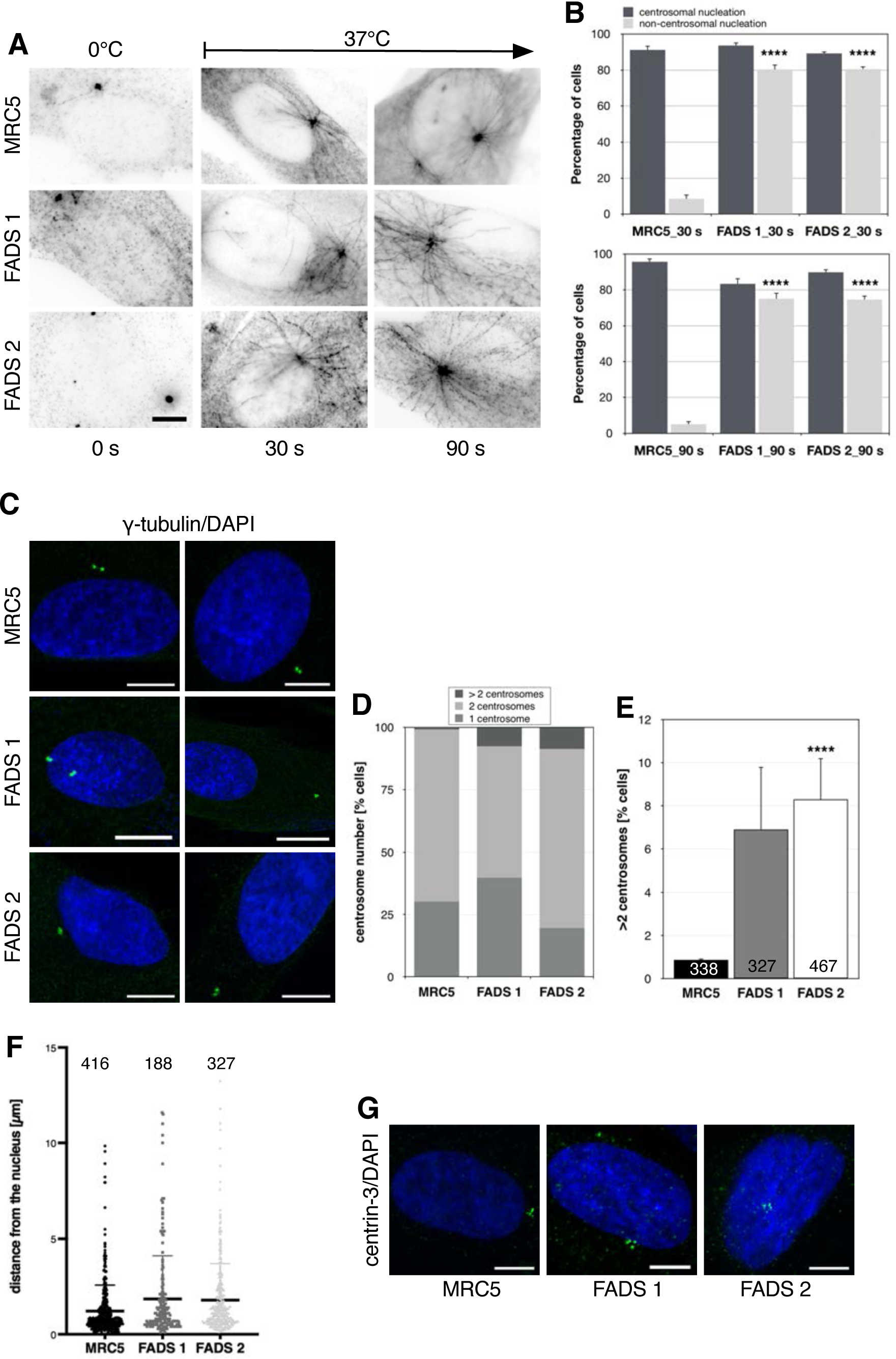
FADS fibroblasts exhibit centrosome abnormalities. (**A**) Representative anti-β-tubulin immunofluorescence of MRC5 and FADS fibroblasts during microtubule regrowth at 37°C after cold shock-induced microtubule depolymerisation at 0°C. FADS cells show non-centrosomal microtubule polymerisation and altered microtubule growth speed. Inverted images of fluorescence signals are shown. Scale bars, 10 µm. (**B**) Quantitative analysis of centrosomal and non-centrosomal microtubule regrowth after 30 s and 90 s at 37°C. (**C**) Abnormal centrosome number and distance in FADS 1 and FADS 2 fibroblasts as compared to MRC5 cells. Cells were stained with an γ-tubulin antibodies (green). DNA was stained with DAPI (blue). Shown are representative confocal images. Quantitative analysis of centrosome number represented as stacked bar plots (**D**) and bar plots (**E**), and (**F**) their distance to the nucleus represented as scatter plots in the indicated fibroblast cell lines. Measurements were done using Fiji/ImageJ. Data present the mean ±SD of at least three independent experiments. P-value ****<0.0001, t-test, two-tailed. The total number of analysed cells were as indicated. (**G**) Centriole numbers for FADS 1 and FADS 2 are similar to those in MRC5 fibroblasts. Cells were stained with anti-centrin-3 antibodies (green). DNA was stained with DAPI (blue). Shown are representative confocal images. Scale bars, 5 µm.

The centrosome is generally the main microtubule organising centre (MTOC) and is responsible for nucleating microtubules (Petry and Vale, 2015). Due to the ncMT nucleation observed in FADS fibroblasts, we next analysed their centrosome number and position. Staining of fibroblasts with γ-tubulin antibodies (Fig. 3C) revealed that FADS fibroblasts have abnormal centrosome numbers, i.e. more than two (Fig. 3D, E). Whereas only about 1% of MRC5 cells had more than two centrosomes, in FADS fibroblasts the percentage increased to 7-8%. Abnormal centrosome number in FADS 1 and FADS 2 did not coincide with an alteration in cell cycle distribution (Fig. 2E). Moreover, centrosomes in FADS cells were more frequently found at larger distance from the nucleus in comparison to control cells (Fig. 3F). Immunostaining with centrin-3 antibodies revealed that centriole number in FADS fibroblasts, however, appeared normal, i.e. each centrosome is comprised of two centrioles (Fig. 3G).

### Ciliary abnormalities in FADS fibroblasts

Centrosome abnormalities are frequently associated with human disease, not only cancer, but also ciliopathies (Bettencourt-Dias et al., 2011). Considering the pleiotropic developmental abnormalities that characterises fetal akinesia, we explored whether defects in primary cilia (PC) formation may contribute to the disease. To address the question, MRC5 control and FADS fibroblasts were subjected to serum-starvation and PC formation was analysed by immunofluorescence microscopy using antibodies against acetylated α-tubulin and Arl13b (ADP ribosylation factor like GTPase 13b), a ciliary marker protein. By both, α 4B) immunostaining, we found that PC were significantly shorter in FADS fibroblasts compared to control cells, and the percentage of ciliated cells was smaller (Fig. 4A, B). Similarly, we observed PC defects in FADS fibroblasts grown on crossbow-shaped micropatterns and stained for acetylated and detyrosinated α tubulin (Fig. S2A).The reduced PC length appears to arise from defects in cilia growth, as PC resorption in FADS fibroblasts was similar to control cells, i.e. not enhanced (Fig. S2B). In contrast to FADS, HGPS and other laminopathies did not coincide with PC defects (Fig. S2C). Expression of wild-type Venus 1-tagged rapsyn (Fig. 4C) or FLAG-tagged rapsyn (Fig. 4D) in FADS 2 cells, which harbour the rapsyn E162K mutant, rescued the ciliary length defect in these cells, in contrast to mutant E162K rapsyn and the respective empty vectors, confirming that the FADS-causing mutation in *RAPSN* contributes to the ciliary defects.

**Figure 4:**
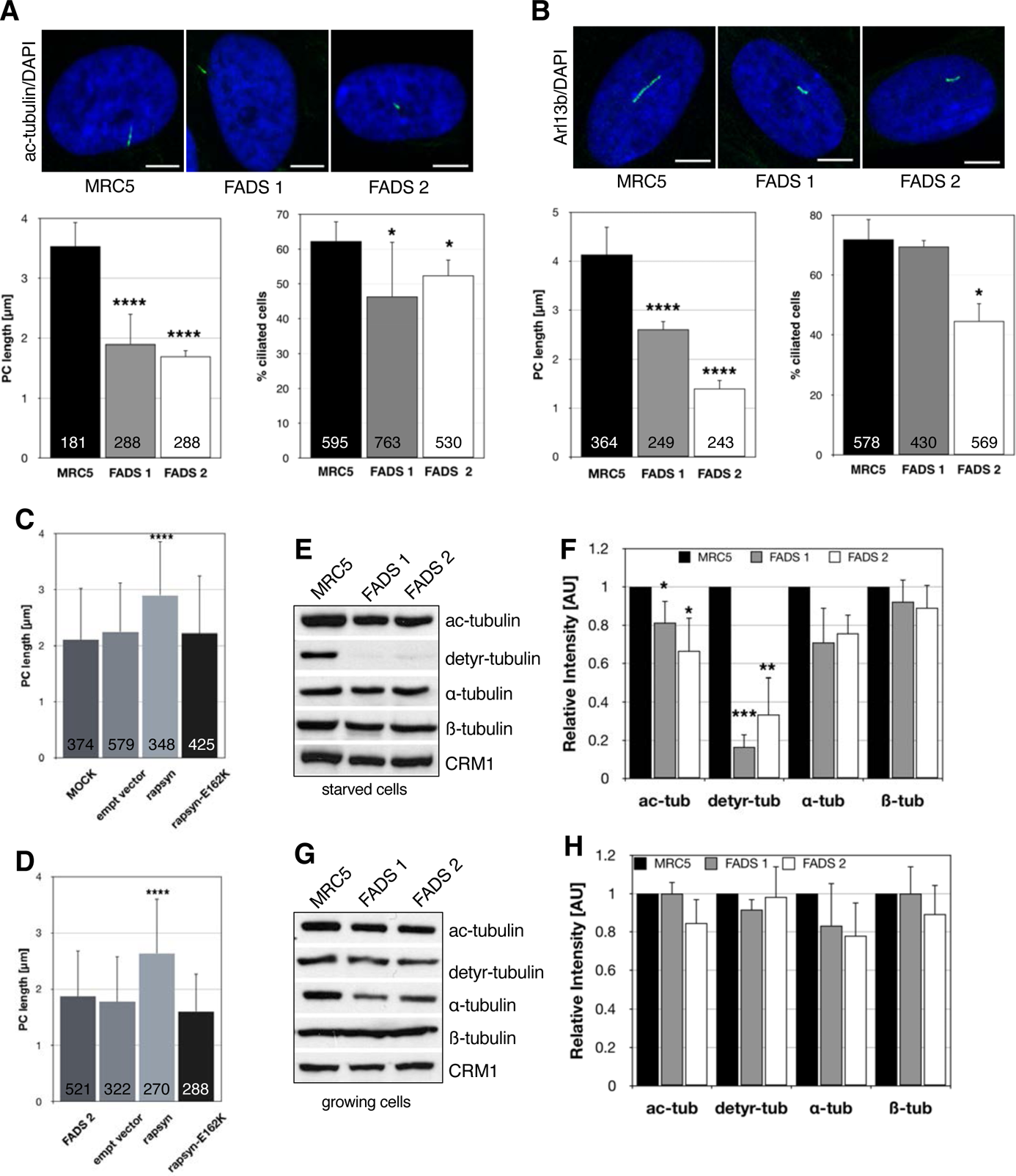
Ciliogenesis is impaired in FADS fibroblasts. Representative confocal images of (**A**) anti-acetylated α-tubulin (ac-tubulin, green) and (**B**) anti-Arl13b (green) immunofluorescence staining in MRC5, FADS1, and FADS2 fibroblasts alongside with quantification of primary cilia (PC) length and percentage of ciliated cells are shown. Fibroblasts were subjected to serum-starvation 48 h before fixation and staining. Expression of Venus1-tagged rapsyn (**C**) or FLAG-tagged rapsyn (**D**) led to a rescue of PC length in FADS 2. PC number was counted manually and PC length was measured using Fiji/ImageJ. Data present the mean ±SD of at least three independent experiments. Cell lysates from (**E**) serum-starved and (**G**) growing fibroblasts were subjected to Western blot analysis using antibodies against acetylated (ac-) α-tubulin, detyrosinated (detyr) α-tubulin, α-tubulin and β-tubulin. Anti-CRM1 antibodies were used as loading control. Relative intensity of acetylated α-tubulin (ac-tub), detyrosinated α-tubulin (detyr-tub), α-tubulin and β-tubulin levels in (**F**) starved and (**H**) growing fibroblasts as determined by densitometry from three independent experiments using Fiji/ImageJ. Total number of analysed cells is indicated at the bottom of each bar. P-values ****<0.0001, ***<0.001, **<0.01, *<0.05, t-test, two-tailed. Scale bars, 10 µm.

PC growth and length is regulated by changes in post-translational modifications (PTMs) of α-tubulin, in particular acetylation, detyrosination, and glycylation of α and β-tubulin (Gadadhar et al., 2017; Gaertig and Wloga, 2008). The levels of detyrosination increase during PC assembly (Gaertig and Wloga, 2008), consistent with assembly defects, we found by immunoblotting significantly lower levels of detyrosinated α-tubulin in serum-starved FADS fibroblasts as compared to serum-starved MRC5 cells (Fig. 4E, F). Also, acetylated α-tubulin was reduced, while the levels of overall α-tubulin and β-tubulin were comparable in FADS and MRC5 control cells (Fig. 4E, F). Variations in α-tubulin PTMs and overall tubulin levels in growing (i.e. non-starved) cells were not significant (Fig. 4G, H), whereas the cellular distribution of detyrosinated α-tubulin in FADS cells appeared more aggregated in immunofluorescence staining (Fig. S2D). Due to a lack of appropriate antibodies against glycylated tubulin, we could not investigate glycylation of α- and β-tubulin in our fibroblasts.

### Depletion of FADS proteins causes ciliary defects

To support the idea that FADS in more general is a ciliopathic disease, we next depleted rapsyn or NUP88 from MRC5 fibroblasts by RNA interference. Mutations in either *RAPSN* or *NUP88* are known genetic causes for FADS (Bonnin et al., 2018; Michalk et al., 2008; Vogt et al., 2008; Winters et al., 2017). Forty-eight hours after siRNA transfection, cells were subjected to serum-starvation to initiate cell cycle exit and PC formation. We found that PCs were significantly shorter upon the respective depletion of rapsyn and NUP88 from MRC5 cells, as revealed by acetylated α-tubulin (Fig. 5A, B) and Arl13b (Fig. 5A, C) staining and immunofluorescence microscopy. Similarly, siRNA-mediated depletion of MuSK, another key factor in the pathogenesis of FADS, resulted in shorter PC (Fig. S3A). Depletion levels of rapsyn and NUP88 were determined by Western blot and bands quantified by densitometry (Fig. S3B-D).

**Figure 5:**
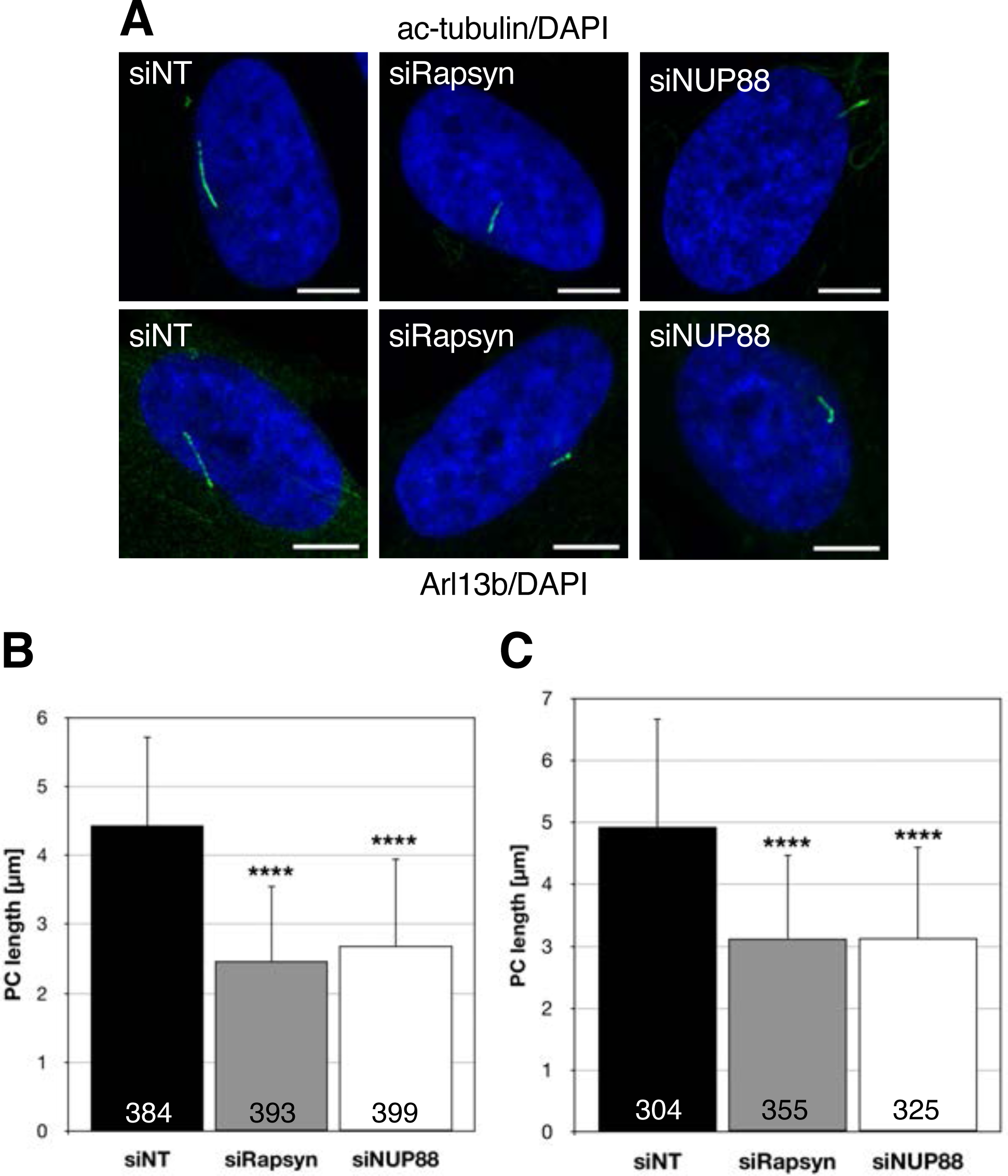
FADS related proteins are critical for ciliogenesis. (**A**) Effects of control (siNT), rapsyn (siRapsyn) or NUP88 (siNUP88) depletion in MRC5 cells subjected 48 h to serum starvation and ciliogenesis. Representative confocal images of cilia marker acetylated α-tubulin (ac-tub; green) and Arl13b (green), and DNA (blue) are shown. The respective quantification of primary cilia (PC) length are shown in (**B)** acetylated α-tubulin and (**C)** Arl13b. Measurements were done using Fiji/ImageJ. Data present the mean ±SD of at least three independent experiments. Total number of analysed cells is indicated at the bottom of each bar. P-values ****<0.0001, t-test, two-tailed. Scale bars, 10 µm.

### FADS proteins localise to centrosomes and cilia

We next determined the intracellular localisation of rapsyn in MRC5 and FADS fibroblasts. As shown in Fig. 6A, rapsyn is distributed throughout the cytoplasm and enriched at centrosomes in all three cell lines. To confirm the association of rapsyn with centrosomes, we performed co-localisation experiments and found that rapsyn indeed co-localised with γ-tubulin at centrosomes in methanol-fixed MRC5 fibroblasts (Fig. 6B). The association of rapsyn with centrosomes was additionally confirmed in the human skeletal muscle precursor rhabdomyosarcoma cell line RH30 (Fig. 6B). Moreover, also NUP88 localised to centrosomes in MRC5 and RH30 cells (Fig. 6B), in contrast to its partner nucleoporins NUP214 and NUP62, as well as NUP93 and NUP153 (Fig. S4A).

**Figure 6:**
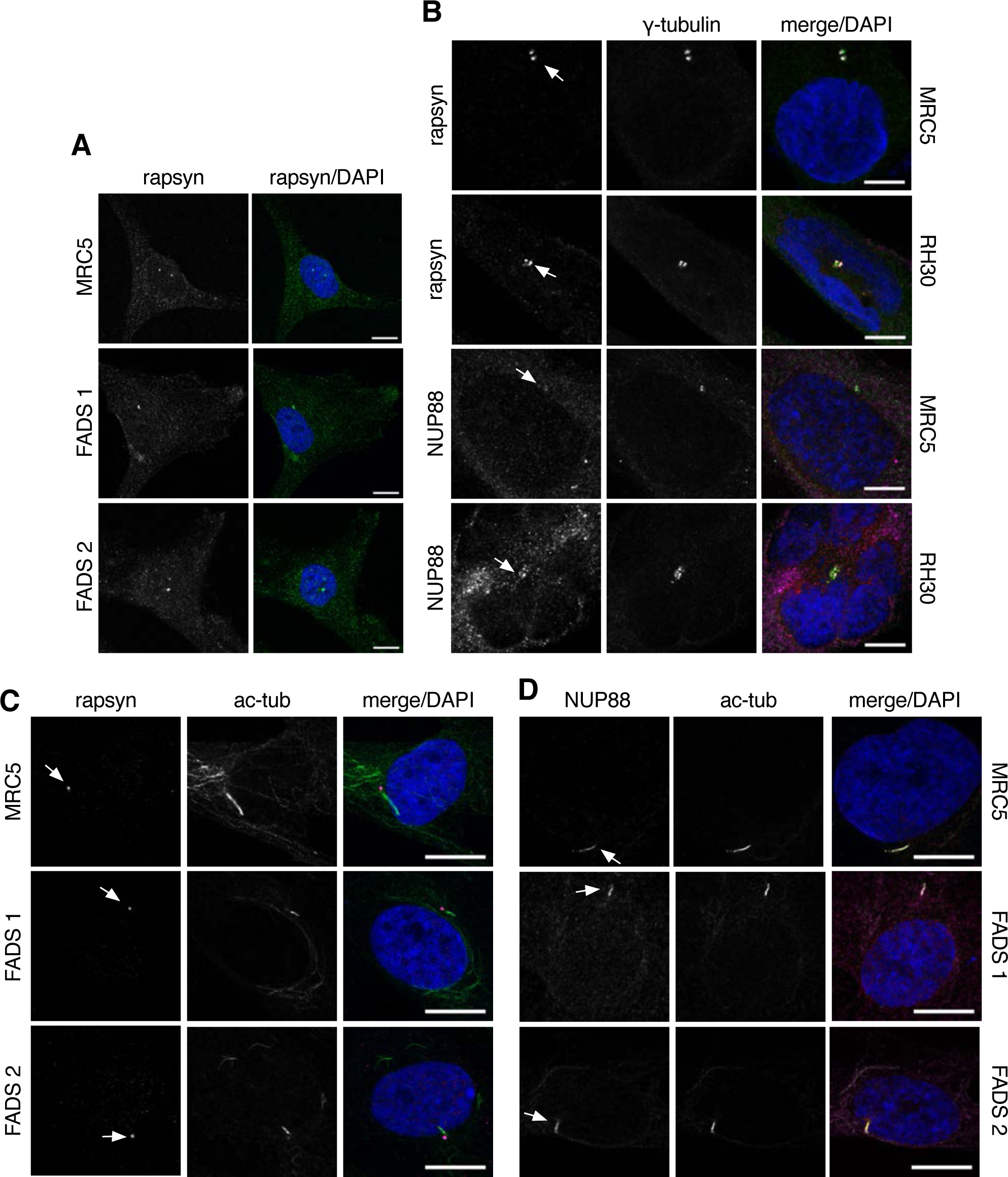
Localisation of rapsyn and NUP88 at centrosomes and primary cilia. (**A**) MRC5 and FADS cells were immunostained with anti-rapsyn antibodies (green) to reveal its intracellular localisation. (**B**) MRC5 and RH30 cells were labelled with anti-rapsyn (magenta) or anti-NUP88 antibodies (magenta), and co-immunostained with anti-γ-tubulin antibodies (green). (**C**) Rapsyn (magenta) and (**D**) NUP88 (magenta) localise to the primary cilium in sub-confluent MRC5, FADS1, and FADS 2 fibroblasts that were serum starved for 48 h. Cilia were visualised by anti-acetylated α-tubulin staining (ac-tub; green). DNA was visualised by DAPI (blue). Rapsyn is located at the cilia base, whereas NUP88 localised along the axoneme of the PC. Shown are representative confocal images. Arrows depict the distinct localisation of the proteins. Scale bars, 10 µm.

In view of the strong link between centrosomes and PC (Chavali et al., 2014; Gupta et al., 2015), we examined the localisation of rapsyn and NUP88 in serum-starved, ciliated MRC5 and FADS fibroblasts. In MRC5 cells, we detected rapsyn at the cilia base (Fig. 6C) and NUP88 along the axoneme of the PC (Fig. 6D), similar to its partner nucleoporins NUP214 and NUP62, but in contrast to the NUP93 and NUP98 (Fig. S4B). Recruitment of rapsyn and NUP88 to the PC was not affected in FADS 1 and FADS 2 fibroblasts, despite the homozygous point mutation of *RAPSN* (c.484G>A) in FADS 2 cells (Fig. 6C, D; Table S1). Besides rapsyn and NUP88, we also detected the FADS protein DOK7 at axoneme in MRC5 and FADS fibroblasts (Fig. S4C), in contrast to MuSK, which did not locate to the cilia base or to axoneme (Fig. S4D).

To further support the link between rapsyn and NUP88 on the one hand and centrosomes and the PC on the other, we next executed proximity ligation assays (PLAs). By PLA, proteins adjacent to the protein of interest with a maximum distance of 40 nm can be detected as PLA amplification foci in the cell (Duolink® PLA Troubleshooting Guide). As shown in Fig. 7A, PLA foci in the cytoplasm of MRC5 fibroblasts were detected for rapsyn with β-tubulin and γ-tubulin, as well as with NUP88. Only a few PLA foci for rapsyn and α-tubulin were detected. The number of rapsyn/NUP88 PLA foci per cell was significantly reduced in FADS 1 and FADS 2 fibroblasts (Fig. 7A, B). Moreover, in FADS 2, rapsyn/β-tubulin PLA foci were almost lacking, whereas their number per cell was increased in FADS 1 (Fig. 7A, B). The amount of rapsyn/γ-tubulin PLA foci per cell was similar to MRC5 in FADS fibroblasts (Fig. 7B).

**Figure 7:**
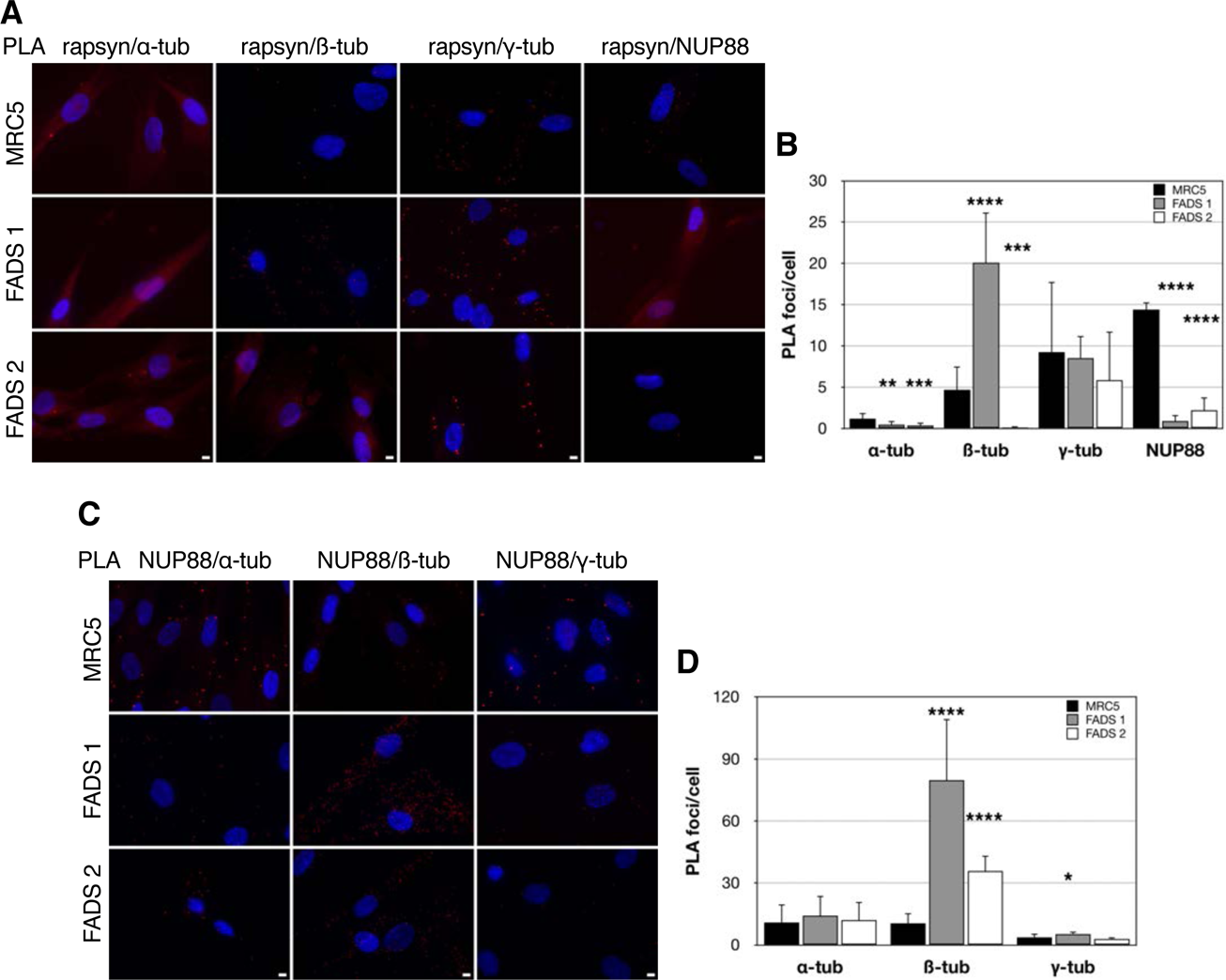
Proximity ligation assays in MRC5, FADS 1 and FADS 2 fibroblasts. (**A**) The close proximity (∼40 nm) between rapsyn and α tubulin (-tub), β-tubulin (β tub), γ-tubulin (γ-tub) or NUP88 was visualised by PLA foci (red). DNA was stained with DAPI (blue). (**B**) Quantification of PLA foci/cell between rapsyn and designated proteins using Fiji/ImageJ. (**C**) The close proximity between NUP88 and -tubulin (-α-tub)or γ-tubulin (γ-tub) was visualised by PLA foci (red). DNA was stained with DAPI (blue). Shown are representative immunofluorescence images. (**D**) Quantification of PLA foci/cell between NUP88 and designated proteins using Fiji/ImageJ. P-values ****<0.0001, ***<0.001, **<0.01, *<0.05, t-test, two-tailed. Scale bars, 10 µm.

Additionally, PLA foci in MRC5 were detected for NUP88 and all three tubulin isoforms (Fig. 7C). The number of PLA foci per cell for NUP88/β-tubulin was significantly increased in FADS 1 and FADS 2, whereas the amount of NUP88/α-tubulin PLA foci per cell was similar in all three fibroblast cell lines (Fig. 7D). In FADS 2, the number of NUP88/ γ-tubulin PLA signals per cell was similar to MRC5, whereas the number was slightly increased in FADS 1 (Fig. 7D). A schematic summary of the PLA results is illustrated in Fig. 8A and control PLAs are presented in Fig. S5A.

**Figure 8:**
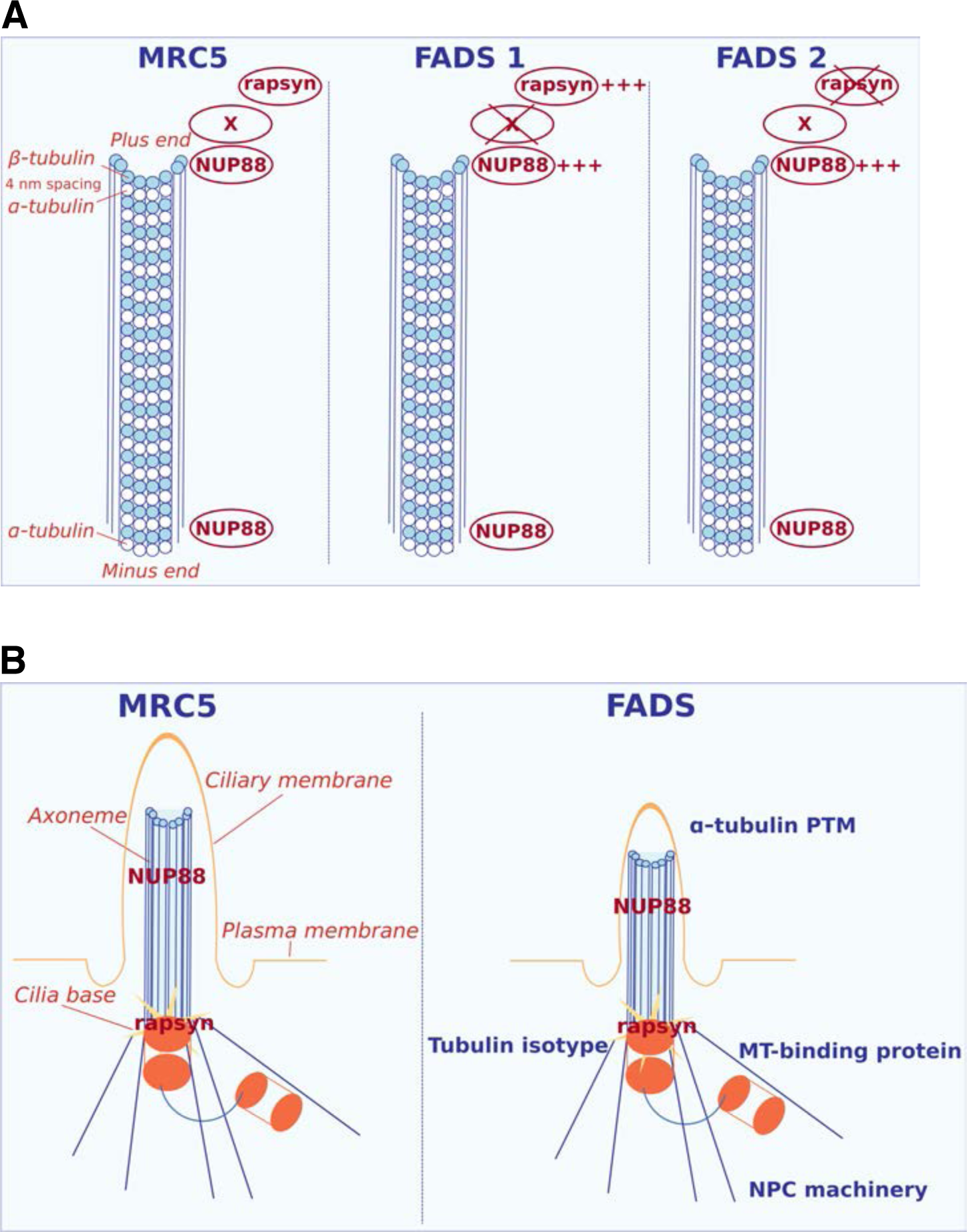
Schematic representations summarising the association of rapsyn and NUP88 with microtubules and primary cilia in control and FADS cells. (**A**) Summary of PLA results in MRC5, FADS 1 and FADS 2 cells (Figure 7). The vicinity of rapsyn and NUP88 at the growing microtubule plus end is abolished in both FADS cell lines. In FADS 2, likely due to the displacement of mutant rapsyn from the microtubule plus end. In FADS 1, possibly due to a mutation (i.e. the unknown disease causing mutation) in an unknown partner (protein X). In both FADS cell lines abolished localisation at the microtubule plus end leads to a possible compensating mechanism of associated proteins (marked with +++). (**B**) Possible mechanisms leading to impaired ciliogenesis in FADS.

## Discussion

FADS is a clinically and genetically heterogeneous constellation, which is characterised by reduced or lack of foetal movement *in utero*. While in the past few years, due to the increased availability of next generation sequencing, significant advances have been made in understanding the molecular aetiology of FADS, the cellular consequences of FADS-related mutations have previously remained largely unstudied. Here we provide evidence that FADS coincides with defects in ciliogenesis, which may account for the pleiotropic morphological defects seen in FADS individuals. We show that primary cilia (PC) length is reduced in fibroblasts derived from two FADS individuals (Fig. 4) and that the respective depletion of the FADS proteins rapsyn, NUP88 and MuSK from control fibroblasts is similarly resulting in shorter PC (Fig. 5, Fig. S3A). Hence, ciliopathic disease underlies the pathogenesis of FADS, at least in a subset of patients.

### FADS fibroblasts have shorter primary cilia

Important for the regulation of PC length are tubulin post-translational modifications (PTMs). PTMs of α-tubulin have been linked to a variety of human pathogenic phenotypes (Magiera et al., 2018). Axonemal microtubules (MTs) are a common target for several PTMs, and the amount of PTMs increases during maturation of the cilium, distinguishing mature cilia from assembling/immature cilia (Wloga et al., 2017). Stable MTs are marked by acetylation, and axonemal MTs are a common target for acetylation contributing to ciliogenesis and ciliary mechano-sensing (Aguilar et al., 2014; Shida et al., 2010). Detyrosinated α-tubulin is enriched at the B-tubules (Johnson, 1998). Since the anterograde intraflagellar transport (IFT) moves along B-tubules, detyrosination has been suggested to stimulate anterograde IFT (Sirajuddin et al., 2014). The ‘tubulin code’, which defines, besides α-tubulin PTMs, also the unique composition of the different tubulin isotypes, is a crucial characteristic of axonemal MTs (Verhey and Gaertig, 2007). Indeed, it has been shown in *C. elegans*, that the loss of the α-tubulin isotype leads to abnormal axoneme ultrastructure and impaired cilia transport (Silva et al., 2017). We hypothesise that a distinct alteration in the ‘tubulin code’, including PTMs, in FADS cells adds to the observed PC defects and probably results in a decreased amount of mature cilia (Fig. 4, Fig. 5, Fig. 8B). We consider indeed defects in anterograde IFT and PC maturation as causative for the reduced PC length and number in FADS, as resorption of PC is similar in FADS and control fibroblasts (Fig. S2B), indicating a regular retrograde IFT. Axonemal MTs can be seen as extensions of the cytoplasmic MT network, which plays an important role in the regulation of cilia by maintaining the balance of available tubulin molecules (Sharma et al., 2011). We are currently examining how the cytoplasmic MT network is organised in FADS cells and can contribute to PC impairments.

### The role of rapsyn and NUP88 in ciliogenesis

How do rapsyn and NUP88 come into the game? A striking connection between nucleoporins, the nucleocytoplasmic transport machinery and cilia is known since quite some years, but the function of nucleoporins at cilia has remained largely obscure (for review see: (Johnson and Malicki, 2019)). A proteomic screen has recently identified NUP88 as part of the cilia proteome (Boldt et al., 2016). Consistent with the localisation of NUP88 to the axoneme of ciliated cells (Fig. 6D), NUP88 appears to be part of the Bardet-Biedl Syndrome (BBSome) coat complex, in complex with DYNLT1 and the WDR19/IFT-A complex (Boldt et al., 2016). Similar to NUP88, we also detected its complex partners NUP214 and NUP62 at the axoneme, in contrast to other nucleoporins, such as NUP98 and NUP93 (Fig. S4B). In contrast to our data presented here, NUP214 and NUP62 have both been previously localised to the cilia base and transition zone (Kee et al., 2012; Takao et al., 2017). This discrepancy might arise from two obvious differences: we examined the endogenous proteins by employing antibodies, whereas in previous studies epitope-tagged versions of NUP214 and NUP62 were analysed. Alternatively, species-specific differences between human (in our study) and rodent (rat or mice in the previous studies) cells/cilia may play a role. Further studies are required to untangle this inconsistency.

NUP88 together with NUP214 and NUP62 has crucial roles in nuclear export by interacting with nuclear export receptors (Bernad et al., 2006; Hutten and Kehlenbach, 2006). Briefly, cargoes are carried from the nucleus to the cytoplasm by nuclear export receptors regulated by the small GTPase Ran. There is emerging evidence that the nuclear export receptor CRM1 regulates non-centrosomal (nc) MT nucleation in a RanGTP-dependent mechanism, and that depletion of CRM1 in yeasts results in increased ncMT nucleation after cold-treatment (Bao et al., 2018; Huang and Avasthi, 2019). Furthermore, Gle1, which has a crucial role in mRNA export and is closely associated with the nuclear pore complex, has been shown to locate to centrosomes and the basal body, and to regulate MT organisation (Jao et al., 2017). It is tempting to speculate that the increased levels of random ncMT nucleation after cold-treatment in FADS cells (Fig. 3A) are caused by similar mechanisms, and thus impair PC formation (Fig. 8B), alter mitotic spindles (Fig. S1A), and concomitantly result in the lower proliferation rates of FADS fibroblasts (Fig. 2). It will be interesting to see how NUP88 is recruited to centrosomes and MTs, and whether this involves similar mechanisms as described for the muscle-specific α-isoform of nuclear envelope protein Nesprin1/SYNE1 (Potter et al., 2018; Potter et al., 2017). Of note, mutations in *SYNE1* have also been described in FADS cases (Attali et al., 2009).

With respect to NUP88 and its association with the BBSome, it has previously been shown, in *C. elegans* and mice, that defects in BBSome components result in defective IFT, which is important for cilium growth, and morphological defects of the cilia (Blacque et al., 2004; Eguether et al., 2014; Keeling et al., 2016; Ou et al., 2007; Rosenbaum and Witman, 2002). This suggests that NUP88 may play a role in IFT as well, and that its loss consequently impairs IFT and cilia growth. Future mechanistic studies should eventually provide more insights in this regard. Interestingly, mutations in WDR19, a binding partner of NUP88 in the BBSome (Boldt et al., 2016), are associated with cranioectodermal dysplasia 4 (CED4), a disorder primarily characterised by craniofacial, skeletal and ectodermal abnormalities (Bredrup et al., 2011).

Rapsyn has thus far not been associated with the cilium. The fact that expression of wild-type rapsyn in the FADS 2 fibroblasts rescued the ciliary defects (Fig. 4C, D), as well as rapsyn’s localisation at centrosomes (Fig. 6A, B) and at the cilia base (Fig. 6C), however, strongly supports the idea that rapsyn plays a role in ciliogenesis. This notion is further strengthened by our PLA assays: firstly, the respective association between rapsyn and β-tubulin and γ-tubulin in PLA assays (Fig. 7A) and, secondly, the disruption of the rapsyn/β-tubulin association in FADS 2 fibroblasts correlates with impaired ciliogenesis in these cells (Fig. 4, Fig. 7A). Interestingly, MACF1 (microtubule-actin crosslinking factor) has been recently identified to bind to AChRs in a rapsyn-dependent manner (Oury et al., 2019). MACF1 is a MT plus end-binding protein shown to promote persistent MT growth and to connect the actin and MT cytoskeletal network (Leung et al., 1999). This novel interaction of rapsyn with MACF1 can be the link to its role in ciliogenesis (Fig. 8B).

Interestingly, our PLA results also suggest a vicinity of rapsyn and NUP88, and β-tubulin/growing MT plus end (only a few PLA signals were observed for rapsyn/α-tubulin). The rapsyn and NUP88 association is abolished in FADS cells (Fig. 7A, B, and Fig. 8A), which in FADS 2 is likely due to the disrupted interface between the mutant rapsyn and β-tubulin, despite augmented association between NUP88 and β-tubulin (Fig. 7C, D) The liaison of both rapsyn and NUP88 to β-tubulin is equally enhanced in FADS 1 cells (Fig. 7A-D), while at the same time rapsyn and NUP88 no longer co-occur at the MT plus end (Fig. 7A, B, and 8A). The underlying molecular basis for this perturbation remains to be elucidated (Fig. 7A, B, and 8A). Another yet to be identified linking protein may account for this effect (Fig. 8A).

Why is less rapsyn in vicinity to α-tubulin than to β-tubulin? The α- and β-tubulin monomers in the MT are spaced approximately 4 nm apart (Díaz et al., 1994). We therefore assume, as discussed above, that rapsyn is predominantly found in vicinity of the growing MT plus end (Fig. 8A), and that its association with α-tubulin cannot be efficiently detected because of the methodological resolution. NUP88, on the contrary, likely also associates with the α-tubulin/shrinking MT minus end (Fig. 8A), and this liaison remains unaffected in both FADS fibroblasts (Fig. 7C,D, and 8A). We hypothesise that the perturbation of FADS proteins at the growing MT plus end interferes with MT dynamics by destabilising and depolymerising MTs. The PLA results at the MT level cannot be directly translated to PC, since further mechanism are regulating ciliogenesis, nevertheless they are hinting to an altered interface between axonemal MTs and rapsyn and NUP88, respectively. Future studies will be required to unravel the underlying molecular mechanisms as to how rapsyn and NUP88 have an impact on cilia growth.

Taken together, our data presented here strongly support the notion that impaired ciliogenesis can underlay FADS and that this contributes to the pleiotropic clinical features that coincide with the disease, in particular to craniofacial and skeletal defects. Moreover, PC are required for proper myoblast differentiation (Fu et al., 2014), which suggests that also the muscular and movement defects seen in FADS individuals at least in part arise from impaired PC formation and growth.

## Material and Methods

All experiments were carried out at room temperature, in duplicate and repeated at least three times unless otherwise stated.

### Plasmids

The pcDNA3.1-rapsyn-FLAG construct was purchased from Genomics Online (ABIN4924584; Aachen, Germany). Rapsyn-Venus1 was generated by Gateway cloning using pDONR223-RAPSN (ABIN5316678; Genomics Online) as donor and pDEST-ORF-V1 (Addgene plasmid #73637, a gift from Darren Saunders; (Croucher et al., 2016)) as destination vector. The presence and the integrity of the *RAPSN* insert was confirmed by sequencing.

Rapsyn-FLAG and rapsyn-Venus1 E162K mutants were generated by site-directed mutagenesis using the QuikChange Lightning site-directed mutagenesis kit (Agilent Technologies, CA, USA) following the manufacturer’s instructions. The primers were: forward 5’-CACACGCGGCACTTGAGCATGGCGTCA-3’, reverse 5’-TGACGCCATGCTCAAGTGCCGCGTGTG-3’. All constructs were verified by DNA sequencing.

### Cell culture and transfection

MRC5 (catalogue number AG05965-G), FADS (FADS 1; catalogue number GM11328), and HGPS (catalogue number AG 01972) fibroblasts were obtained from Coriell (Human Genetic Cell Repository; Coriell Institute, Hamden, NJ, USA). FADS 2 fibroblasts were from an individual harbouring a homozygous c.484G>A (p.Glu162Lys) variant in the *RAPSN* gene (NM_005055.4) as described previously (Winters et al., 2017). FADS 1 cells were grown in Minimum Essential Medium (MEM) Alpha (Lonza, Basel, Switzerland) supplemented with 15% foetal bovine serum (FBS) plus penicillin and streptomycin. HGPS cells were grown in MEM medium (Life Technologies Gibco, Gent, Belgium) supplemented with 15% FBS, non-essential amino acids plus penicillin/streptomycin. MRC5 and FADS 2 fibroblasts were grown in MEM medium supplemented with 10% FBS and penicillin/streptomycin. All cell lines were grown at 37°C in a humidified incubator with 5% CO_2_ atmosphere.

Transfection with siRNAs was carried out using Lipofectamine RNAiMAX (Life Technologies Invitrogen) following the instructions of the manufacturer. siRNAs were from Dharmacon (Lafayette, CO, USA): *RAPSN* (L-006550-00), *MUSK* (L-03158-00), *NUP88* (L-017547-01-0005), and non-targeting siRNAs (D-001810-10). Plasmids were transfected using Lipofectamine 3000 (Life Technologies Invitrogen) following the instructions of the manufacturer.

### EdU cell proliferation assay

For EdU incorporation, cells were plated on coverslips in a 24-well plate and grown to 70-80% confluency. Cells were next incubated for 1 h with medium containing 10 µM EdU, according to the manufacturer’s instructions (Click-iT Edu Alexa Fluor 594 Imaging Kit, Invitrogen). After incubation, cells were fixed for 15 min in 3.7% formaldehyde, washed twice in PBS containing 2% bovine serum albumin (BSA), permeabilised with PBS containing 0.5% Triton X-100 for 20 min, and washed twice in PBS containing 2% BSA. Cells were next incubated for 30 min with Click-iT reaction solution according to the manufacturer’s instructions, washed twice in PBS containing 2% BSA, washed two times in PBS. Coverslips were mounted with a drop of Mowiol-4088 (Sigma-Aldrich, St. Louis, MO, USA) containing DAPI (1 µg ml^-1^; Sigma-Aldrich). Cells were imaged using a Zeiss Observer.Z1 microscope (Zeiss, Oberkochen, Germany). Images were recorded using the microscope system software and processed using Fiji/ImageJ (Version 1.52p; (Schindelin et al., 2012)). EdU positive cells were counted manually.

### Senescence assays

Cells were seeded in 6-well culture dishes at a density of 0.4 ×10^5^ cells/well 48 h prior to the experiment. Cellular senescence was analysed using the Senescence β-Galactosidase Staining Kit (Cell Signalling Technology, Cambridge, UK) according to the protocol from the manufacturer. Development of the blue colour in senescent cells while the β-galactosidase was still on the culture dishes was evaluated using a Zeiss Observer.Z1 microscope. Images were recorded using the microscope system software and processed using Adobe Photoshop (Version 12.0; Adobe Systems, Mountain View, CA, USA). β-galactosidase positive cells were counted manually.

### Cell cycle assays

Fibroblasts were plated in 10 cm dishes (0.6×10^6^ cells per dish), harvested after 24 h by trypsinization, collected with initial culture medium, washed with PBS, and fixed overnight at −20°C with 70% ethanol. For flow cytometric analysis, the cells were washed with PBS and were first incubated for 5 min at room temperature with RNase A (0.2 mg/ml; Invitrogen), followed by a 15 min incubation at 37°C with propidium iodide (PI; Sigma-Aldrich) solution (1 mg ml^-1^ PI, 10% Triton X-100 in PBS). Cells were analysed by flow cytometry, using a FACS CantoII (BD Biosciences). Cell cycle distribution was analysed using the FlowJo software.

### Microtubule assays

In order to assess kinetochore-fibre microtubule stability, cells were either put on ice for 10 min or treated with 10 µM (complete loss) or 170 nM (low polymerisation) of nocodazole (Sigma-Aldrich). For microtubule repolymerisation assays, cells were put on ice for 10 min and then incubated again in a 37°C humidified 5% CO_2_ atmosphere for 0, 30 and 90 s.

Next, cells were fixed with 4% formaldehyde in PBS for 5 min, permeabilised with 0.5% Triton X-100 in PBS for 5 min and then fixed again. Blocking was performed with 2% BSA/0.1% Triton X-100 in PBS for 30 min. All antibodies were diluted in blocking solution. Primary antibodies were incubated at 4°C over-night in a humidified chamber. Secondary antibodies were incubated 1 h in the dark. Excess antibodies after primary and secondary antibody staining were removed by three washing steps using 0.1% Triton X-100 in PBS for 5 min. Cover slips were mounted onto microscope slides with Mowiol-4088 containing DAPI. Cells were imaged using a Zeiss Observer.Z1 microscope. Images were recorded using the microscope system software and processed using Fiji/ImageJ and Adobe Photoshop. Microtubule nucleation was counted manually.

### Doubling time

Cells were seeded in 6-well culture plates at a density of 0.4 x1^5^ cells/well. Cells were counted using a TC20TM Automated Cell Counter (Bio-Rad, California, USA) at indicated time points, over a range of at least 168 h. Only the linear parts of the count data were used in the analysis.

Growth curve analysis (GCA) and growth constant k (slope of linear regression line) and doubling time T_d_ calculation were calculated using the multilevel regression technique using R (version 3.6.0) and R Studio (version 1.2.1335; R: A language and environment for statistical computing. R Foundation for Statistical Computing, Vienna, Austria. https://www.R-project.org) as described earlier (Mirman, 2014). By using GCA, both group-level and individual-level effects on cellular growth were analysed. The linear model (m.1; Fig. 2A) suggests effects of the different cell lines on proliferation and the intercept model (m.0; Fig. S5B) suggests constant differences in proliferation randomly assigned to the different cell lines. Statistical comparison (Pearson’s χ test with two degrees of freedom of the models revealed that the linear regression model m.1 was used as regression model of choice (P < 0.001)). A third model for regression calculation was evaluated, which included unmeasured individual properties affecting cell growth, but this model did not improve the regression calculation (data not shown).

### Immunofluorescence

Cells were grown on coverslips in 24-well plates and fixed either with 2% formaldehyde for 15 min or at −20°C with methanol (for γ-tubulin and centrin-3 staining) for 10 min, washed three times for 10 min with PBS, and permeabilised with PBS containing 1% BSA and 0.2% Triton X-100 for 10 min on ice. Next, the cells were washed three times for 10 min in PBS containing 2% BSA and incubated with the appropriate primary antibodies for 1 h at RT or over-night at 4°C, washed three times in PBS containing 1% BSA and incubated with the appropriate secondary antibodies for 1 h, washed four times 10 min with PBS and mounted with a drop of Mowiol-4088 containing DAPI. Cells were imaged using a Zeiss LSM-710 confocal laser-scanning microscope (Zeiss, Oberkochen, Germany). Images were recorded using the microscope system software and processed using Fiji/ImageJ and Adobe Photoshop.

The following antibodies were used as primary antibodies: monoclonal mouse anti-lamin A/C (1:30; ab40567; Abcam, Cambridge, UK), mouse anti-γ-tubulin (1:1000; ab11316; Abcam), mouse anti-β-tubulin (1:1000; clone KMX-1; Merck Millipore, Massachusetts, USA), mouse anti-acetylated α Aldrich), mouse anti-DOK7 (1:50; NBP2-02073; Novus Biologicals, Abingdon, UK), mouse anti-NUP88 (1:500; 611896; BD Biosciences, San Jose, CA, USA), mouse anti-Ki-67 (1:100; NCL-L-Ki67-MM1; Novocastra Laboratories, Newcastle upon Tyne, UK), and rat an α-tubulin (1:1000; MCA78G; Bio-Rad AbD Serotec), polyclonal rabbit lamin B1 (1:300; ab16048; Abcam), rabbit anti-γ-tubulin (1:100; 15176-1-AP; Proteintech, Mancester, UK), rabbit anti-centrin-3 (1:1000; PA5-35865; Thermo Scientific, Rockford, USA), rabbit anti-acetylated α-tubulin (1:800; 5335; Cell Signalling Technology, Leiden, The Netherlands), rabbit anti-Arl13b (1:200; 17711-1-AP; Proteintech), rabbit anti-rapsyn (1:200; NBP1-85537; Novus Biologicals), rabbit anti-MuSK (1:100; PA5-14705; Thermo Scientific).

Secondary antibodies were the corresponding goat anti-mouse IgG Alexa Fluor 568, goat anti-rabbit IgG Alexa Fluor 568, goat anti-mouse IgG Alexa Fluor 488 and goat anti-rabbit IgG Alexa Fluor 488. All secondary antibodies were diluted 1:1000 and purchased from Invitrogen.

### Western blotting

Cells were lysed in lysis buffer (50 mM Tris-HCl, pH 7.8, 150 mM NaCl, 1% Nonidet-P40 and protease inhibitor cocktail tablets (Roche, Basel, Switzerland)). 30 µg of protein were loaded and separated by sodium dodecyl sulfate-polyacrylamide gel electrophoresis (SDS-PAGE). The proteins were transferred onto a PVDF membrane (Immobilon-P, Merck Millipore) and the membranes were blocked with TBS containing 0.1% Tween 20 and 5% non-fat dry milk for 1 h. The membranes were then incubated over-night at 4°C in blocking solution containing a primary antibody followed by washing 3x in TBS containing 0.1% Tween 20 and 5% non-fat dry. The membranes were next incubated with secondary antibodies for 1 h, washed 3x in TBS containing 0.1% Tween 20 and developed. X-ray films were scanned and processed using Fiji/ImageJ. Gel densitometry measurement was done with Fiji/ImageJ.

The following antibodies were used as primary antibodies: monoclonal mouse anti-lamin A/C (1:200; ab8984, Abcam), mouse anti-NUP88 (1:1000), mouse anti-DOK7 (1:1000), mouse anti-CRM1 (1:1000; 611832, BD Biosciences), and mouse anti-β-tubulin (1:1000).

Polyclonal rabbit anti-lamin B1 (1:300), rabbit anti-rapsyn (1:500), rabbit anti-MuSK (1:1000), rabbit anti-actin (1:1000; A2066, Sigma-Aldrich), rabbit anti-acetylated α-tubulin (1:1000), rabbit anti-detyrosinated α -tubulin (1:1000), and rabbit anti-α-tubulin (1:4000; ab18251, Abcam).

Secondary antibodies were either alkaline phosphatase-coupled IgG antibodies (1:10.000) from Sigma-Aldrich or horseradish peroxidase-coupled IgG antibodies (1:8000) from Cell Signalling Technology.

### Proximity ligation assay

Cells grown on glass cover slips were fixed with 4% PFA in PBS for 5 min or fixed at −20°C with methanol (for γ-tubulin) for 10 min, permeabilised with 0.5% Triton X-100 in PBS for 5 min and then fixed again. Blocking was performed with 2% BSA/0.1% Triton X-100 in PBS for 30 min.

All antibodies used for proximity ligation assay (PLA) were diluted in blocking solution. Antibodies rabbit anti-rapsyn, rat anti-α-tubulin, mouse anti-β-tubulin, rabbit anti-γ-tubulin, mouse anti-NUP88, and rabbit anti-NUP88 (kind gift from Dr. Ulrike Kutay, ETH Zurich, Switzerland) were incubated at 4°C over-night in a humidified chamber. Excess antibodies were removed by three washing steps using 0.1% Triton X-100 in PBS for 5 min. PLA was performed using Duolink® PLA Fluorescent Detection (red) with anti-mouse PLUS and anti-rabbit MINUS oligonucleotides (Sigma-Aldrich). PLA was performed as described elsewhere (Lin et al., 2015). Wash buffers were prepared as follows: wash buffer A (0.01 M Tris, pH 7.4, 0.15 M NaCl; 0.05% Tween 20) and wash buffer B (0.2 M Tris, pH 7.5, 0.1 M NaCl). Cover slips were mounted onto microscope slides with Mowiol-4088 containing DAPI. Cells were imaged using a Zeiss Observer.Z1 microscope. Images were recorded using the microscope system software and processed using Fiji/ImageJ and Adobe Photoshop. PLA foci were counted using a macro written on Fiji/ImageJ: briefly, parameters and scale were set, images thresholded (Median, Gaussian blur, MaxEntropy) and particles (PLA foci) counted.

### Statistical analyses

All plots and statistics (despite GCA models) were generated using GraphPad Prism (Version 8;GraphPad Software Inc., CA, USA) or Apple Numbers (Apple Inc., CA, USA). Two-tailed t-test was performed. During evaluation of the results a confidence interval α of 95% and p values lower than 0.05 were considered as statistically significant.

### Image design

Schematic representations were designed using the open-source software Inkscape 0.91 (by running XQuartz 2.7.11 on macOS). Colours were adapted from http://www.ColorBrewer.org by Cynthia A. Brewer (Brewer).

## Acknowledgements

We are grateful to Drs. Ulrike Kutay (ETH Zürich, Switzerland) and Darren Saunders (UNSW Sydney, Australia) for sharing reagents. Confocal images were acquired at the CMMI, which is supported by the European Regional Development Fund (ERDF).

No competing interests declared.

This work was supported by grants from the Fédération Wallonie-Bruxelles (ARC 4.110.F.000092F).

## References

1. Aguilar, A., Becker, L., Tedeschi, T., Heller, S., Iomini, C. and Nachury, M. V. (2014). Alpha-tubulin K40 acetylation is required for contact inhibition of proliferation and cell-substrate adhesion. Mol Biol Cell 25, 1854–66.

2. Aittaleb, M., Chen, P. J. and Akaaboune, M. (2015). Failure of lysosome clustering and positioning in the juxtanuclear region in cells deficient in rapsyn. J Cell Sci 128, 3744–56.

3. Antolik, C., Catino, D. H., O’Neill, A. M., Resneck, W. G., Ursitti, J. A. and Bloch, R. J. (2007). The actin binding domain of ACF7 binds directly to the tetratricopeptide repeat domains of rapsyn. Neuroscience 145, 56–65.

4. Attali, R., Warwar, N., Israel, A., Gurt, I., McNally, E., Puckelwartz, M., Glick, B., Nevo, Y., Ben-Neriah, Z. and Melki, J. (2009). Mutation of SYNE-1, encoding an essential component of the nuclear lamina, is responsible for autosomal recessive arthrogryposis. Hum Mol Genet 18, 3462–9.

5. Avalos, Y., Pena-Oyarzun, D., Budini, M., Morselli, E. and Criollo, A. (2017). New Roles of the Primary Cilium in Autophagy. Biomed Res Int 2017, 4367019.

6. Bao, X. X., Spanos, C., Kojidani, T., Lynch, E. M., Rappsilber, J., Hiraoka, Y., Haraguchi, T. and Sawin, K. E. (2018). Exportin Crm1 is repurposed as a docking protein to generate microtubule organizing centers at the nuclear pore. Elife 7.

7. Bartoli, M., Ramarao, M. K. and Cohen, J. B. (2001). Interactions of the rapsyn RING-H2 domain with dystroglycan. J Biol Chem 276, 24911–7.

8. Beecroft, S. J., Lombard, M., Mowat, D., McLean, C., Cairns, A., Davis, M., Laing, N. G. and Ravenscroft, G. (2018). Genetics of neuromuscular fetal akinesia in the genomics era. J Med Genet 55, 505–514.

9. Bernad, R., Engelsma, D., Sanderson, H., Pickersgill, H. and Fornerod, M. (2006). Nup214-Nup88 nucleoporin subcomplex is required for CRM1-mediated 60 S preribosomal nuclear export. J Biol Chem 281, 19378–86.

10. Bettencourt-Dias, M., Hildebrandt, F., Pellman, D., Woods, G. and Godinho, S. A. (2011). Centrosomes and cilia in human disease. Trends Genet 27, 307–15.

11. Blacque, O. E., Reardon, M. J., Li, C., McCarthy, J., Mahjoub, M. R., Ansley, S. J., Badano, J. L., Mah, A. K., Beales, P. L., Davidson, W. S. et al. (2004). Loss of C. elegans BBS-7 and BBS-8 protein function results in cilia defects and compromised intraflagellar transport. Genes Dev 18, 1630–42.

12. Boldt, K., van Reeuwijk, J., Lu, Q., Koutroumpas, K., Nguyen, T. M., Texier, Y., van Beersum, S. E., Horn, N., Willer, J. R., Mans, D. A. et al. (2016). An organelle-specific protein landscape identifies novel diseases and molecular mechanisms. Nat Commun 7, 11491.

13. Bonnin, E., Cabochette, P., Filosa, A., Juhlen, R., Komatsuzaki, S., Hezwani, M., Dickmanns, A., Martinelli, V., Vermeersch, M., Supply, L. et al. (2018). Biallelic mutations in nucleoporin NUP88 cause lethal fetal akinesia deformation sequence. PLoS Genet 14, e1007845.

14. Bredrup, C., Saunier, S., Oud, M. M., Fiskerstrand, T., Hoischen, A., Brackman, D., Leh, S. M., Midtbo, M., Filhol, E., Bole-Feysot, C. et al. (2011). Ciliopathies with skeletal anomalies and renal insufficiency due to mutations in the IFT-A gene WDR19. Am J Hum Genet 89, 634–43.

15. Buck, S. B., Bradford, J., Gee, K. R., Agnew, B. J., Clarke, S. T. and Salic, A. (2008). Detection of S-phase cell cycle progression using 5-ethynyl-2’-deoxyuridine incorporation with click chemistry, an alternative to using 5-bromo-2’-deoxyuridine antibodies. Biotechniques 44, 927–9.

16. Burden, S. J. (2002). Building the vertebrate neuromuscular synapse. J Neurobiol 53, 501–11.

17. Burden, S. J., Huijbers, M. G. and Remedio, L. (2018). Fundamental Molecules and Mechanisms for Forming and Maintaining Neuromuscular Synapses. Int J Mol Sci 19.

18. Chavali, P. L., Putz, M. and Gergely, F. (2014). Small organelle, big responsibility: the role of centrosomes in development and disease. Philos Trans R Soc Lond B Biol Sci 369.

19. Croucher, D. R., Iconomou, M., Hastings, J. F., Kennedy, S. P., Han, J. Z., Shearer, R. F., McKenna, J., Wan, A., Lau, J., Aparicio, S. et al. (2016). Bimolecular complementation affinity purification (BiCAP) reveals dimer-specific protein interactions for ERBB2 dimers. Sci Signal 9, ra69.

20. Díaz JF, Pantos E, Bordas J, Andreu JM. (1994). Solution structure of GDP-tubulin double rings to 3 nm resolution and comparison with microtubules. J Mol Biol 238:214–225.

21. Dobbins, G. C., Luo, S., Yang, Z., Xiong, W. C. and Mei, L. (2008). alpha-Actinin interacts with rapsyn in agrin-stimulated AChR clustering. Mol Brain 1, 18.

22. Drozdz, M. M., Jiang, H., Pytowski, L., Grovenor, C. and Vaux, D. J. (2017). Formation of a nucleoplasmic reticulum requires de novo assembly of nascent phospholipids and shows preferential incorporation of nascent lamins. Sci Rep 7, 7454.

23. Eguether, T., San Agustin, J. T., Keady, B. T., Jonassen, J. A., Liang, Y., Francis, R., Tobita, K., Johnson, C. A., Abdelhamed, Z. A., Lo, C. W. et al. (2014). IFT27 links the BBSome to IFT for maintenance of the ciliary signaling compartment. Dev Cell 31, 279–290.

24. Engel, A. G. (2012). Current status of the congenital myasthenic syndromes. Neuromuscul Disord 22, 99–111.

25. Frail, D. E., Musil, L. S., Buonanno, A. and Merlie, J. P. (1989). Expression of RAPsyn (43K protein) and nicotinic acetylcholine receptor genes is not coordinately regulated in mouse muscle. Neuron 2, 1077–86.

26. Fu, W., Asp, P., Canter, B. and Dynlacht, B. D. (2014). Primary cilia control hedgehog signaling during muscle differentiation and are deregulated in rhabdomyosarcoma. Proc Natl Acad Sci U S A 111, 9151–6.

27. Gadadhar, S., Dadi, H., Bodakuntla, S., Schnitzler, A., Bieche, I., Rusconi, F. and Janke, C. (2017). Tubulin glycylation controls primary cilia length. J Cell Biol 216, 2701–2713.

28. Gaertig, J. and Wloga, D. (2008). Ciliary tubulin and its post-translational modifications. Curr Top Dev Biol 85, 83–113.

29. Galati, D. F., Sullivan, K. D., Pham, A. T., Espinosa, J. M. and Pearson, C. G. (2018). Trisomy 21 Represses Cilia Formation and Function. Dev Cell 46, 641–650 e6.

30. Garcia, G3rd.,, Raleigh, D. R. and Reiter, J. F. (2018). How the Ciliary Membrane Is Organized Inside-Out to Communicate Outside-In. Curr Biol 28, R421–R434.

31. Gerdes, J., Lemke, H., Baisch, H., Wacker, H. H., Schwab, U. and Stein, H. (1984). Cell cycle analysis of a cell proliferation-associated human nuclear antigen defined by the monoclonal antibody Ki-67. J Immunol 133, 1710–5.

32. Ghasemizadeh, A., Christin, E., Guiraud, A., Couturier, N., Risson, V., Girard, E., Jagla, C., Soler, C., Laddada, L., Sanchez, C. et al. (2019). Skeletal muscle MACF1 maintains myonuclei and mitochondria localization through microtubules to control muscle functionalities. bioRxiv 636464, doi: https://doi.org/10.1101/636464.

33. Ghazanfari, N., Fernandez, K. J., Murata, Y., Morsch, M., Ngo, S. T., Reddel, S. W., Noakes, P. G. and Phillips, W. D. (2011). Muscle specific kinase: organiser of synaptic membrane domains. Int J Biochem Cell Biol 43, 295–8.

34. Gupta, G. D., Coyaud, E., Goncalves, J., Mojarad, B. A., Liu, Y., Wu, Q., Gheiratmand, L., Comartin, D., Tkach, J. M., Cheung, S. W. et al. (2015). A Dynamic Protein Interaction Landscape of the Human Centrosome-Cilium Interface. Cell 163, 1484–99.

35. Hall, J. G. (2009). Pena-Shokeir phenotype (fetal akinesia deformation sequence) revisited. Birth Defects Res A Clin Mol Teratol 85, 677–94.

36. Huang, S. and Avasthi, P. (2019). RanGTP regulates cilium formation and ciliary trafficking of a kinesin-II subunit independent of its nuclear functions. bioRxiv 562272, doi: https://doi.org/10.1101/562272.

37. Hutten, S. and Kehlenbach, R. H. (2006). Nup214 is required for CRM1-dependent nuclear protein export in vivo. Mol Cell Biol 26, 6772–85.

38. Jao, L. E., Akef, A. and Wente, S. R. (2017). A role for Gle1, a regulator of DEAD-box RNA helicases, at centrosomes and basal bodies. Mol Biol Cell 28, 120–127.

39. Jevtic, P., Schibler, A. C., Wesley, C. C., Pegoraro, G., Misteli, T. and Levy, D. L. (2019). The nucleoporin ELYS regulates nuclear size by controlling NPC number and nuclear import capacity. EMBO Rep 20.

40. Johnson, C. A. and Malicki, J. J. (2019). The Nuclear Arsenal of Cilia. Dev Cell 49, 161–170.

41. Johnson, K. A. (1998). The axonemal microtubules of the Chlamydomonas flagellum differ in tubulin isoform content. J Cell Sci 111 (Pt 3), 313–20.

42. Kee, H. L., Dishinger, J. F., Blasius, T. L., Liu, C. J., Margolis, B. and Verhey, K. J. (2012). A size-exclusion permeability barrier and nucleoporins characterize a ciliary pore complex that regulates transport into cilia. Nat Cell Biol 14, 431–7.

43. Keeling, J., Tsiokas, L. and Maskey, D. (2016). Cellular Mechanisms of Ciliary Length Control. Cells 5.

44. Kim, N. and Burden, S. J. (2008). MuSK controls where motor axons grow and form synapses. Nat Neurosci 11, 19–27.

45. Kim, N., Stiegler, A. L., Cameron, T. O., Hallock, P. T., Gomez, A. M., Huang, J. H., Hubbard, S. R., Dustin, M. L. and Burden, S. J. (2008). Lrp4 is a receptor for Agrin and forms a complex with MuSK. Cell 135, 334–42.

46. Leung, C. L., Sun, D., Zheng, M., Knowles, D. R. and Liem, R. K. (1999). Microtubule actin cross-linking factor (MACF): a hybrid of dystonin and dystrophin that can interact with the actin and microtubule cytoskeletons. J Cell Biol 147, 1275–86.

47. Li, L., Cao, Y., Wu, H., Ye, X., Zhu, Z., Xing, G., Shen, C., Barik, A., Zhang, B., Xie, X. et al. (2016). Enzymatic Activity of the Scaffold Protein Rapsyn for Synapse Formation. Neuron 92, 1007–1019.

48. Li, N., Qiao, C., Lv, Y., Yang, T., Liu, H., Yu, W. Q. and Liu, C. X. (2019). Compound heterozygous mutation of MUSK causing fetal akinesia deformation sequence syndrome: A case report. World J Clin Cases 7, 3655–3661.

49. Lin, M. Z., Martin, J. L. and Baxter, R. C. (2015). Proximity Ligation Assay (PLA) to Detect Protein-protein Interactions in Breast Cancer Cells. Bio-protocol 5, e1479.

50. Luiskandl, S., Woller, B., Schlauf, M., Schmid, J. A. and Herbst, R. (2013). Endosomal trafficking of the receptor tyrosine kinase MuSK proceeds via clathrin-dependent pathways, Arf6 and actin. FEBS J 280, 3281–97.

51. Luo, W., Ruba, A., Takao, D., Zweifel, L. P., Lim, R. Y. H., Verhey, K. J. and Yang, W. (2017). Axonemal Lumen Dominates Cytosolic Protein Diffusion inside the Primary Cilium. Sci Rep 7, 15793.

52. Magiera, M. M., Singh, P., Gadadhar, S. and Janke, C. (2018). Tubulin Posttranslational Modifications and Emerging Links to Human Disease. Cell 173, 1323–1327.

53. Marchand, S., Bignami, F., Stetzkowski-Marden, F. and Cartaud, J. (2000). The myristoylated protein rapsyn is cotargeted with the nicotinic acetylcholine receptor to the postsynaptic membrane via the exocytic pathway. J Neurosci 20, 521–8.

54. Marchand, S., Devillers-Thiery, A., Pons, S., Changeux, J. P. and Cartaud, J. (2002). Rapsyn escorts the nicotinic acetylcholine receptor along the exocytic pathway via association with lipid rafts. J Neurosci 22, 8891–901.

55. Mazhar, S. and Herbst, R. (2012). The formation of complex acetylcholine receptor clusters requires MuSK kinase activity and structural information from the MuSK extracellular domain. Mol Cell Neurosci 49, 475–86.

56. Michalk, A., Stricker, S., Becker, J., Rupps, R., Pantzar, T., Miertus, J., Botta, G., Naretto, V. G., Janetzki, C., Yaqoob, N. et al. (2008). Acetylcholine receptor pathway mutations explain various fetal akinesia deformation sequence disorders. Am J Hum Genet 82, 464–76.

57. Mihailovska, E., Raith, M., Valencia, R. G., Fischer, I., Al Banchaabouchi, M., Herbst, R. and Wiche, G. (2014). Neuromuscular synapse integrity requires linkage of acetylcholine receptors to postsynaptic intermediate filament networks via rapsyn-plectin 1f complexes. Mol Biol Cell 25, 4130–49.

58. Mirman, D. (2014). Growth curve analysis and visualization using R. New York: Chapman & Hall/CRC The R Series.

59. Mirvis, M., Stearns, T. and James Nelson, W. (2018). Cilium structure, assembly, and disassembly regulated by the cytoskeleton. Biochem J 475, 2329–2353.

60. Moransard, M., Borges, L. S., Willmann, R., Marangi, P. A., Brenner, H. R., Ferns, M. J. and Fuhrer, C. (2003). Agrin regulates rapsyn interaction with surface acetylcholine receptors, and this underlies cytoskeletal anchoring and clustering. J Biol Chem 278, 7350–9.

61. Musil, L. S., Frail, D. E. and Merlie, J. P. (1989). The mammalian 43-kD acetylcholine receptor-associated protein (RAPsyn) is expressed in some nonmuscle cells. J Cell Biol 108, 1833–40.

62. Okada, K., Inoue, A., Okada, M., Murata, Y., Kakuta, S., Jigami, T., Kubo, S., Shiraishi, H., Eguchi, K., Motomura, M. et al. (2006). The muscle protein Dok-7 is essential for neuromuscular synaptogenesis. Science 312, 1802–5.

63. Ou, G., Koga, M., Blacque, O. E., Murayama, T., Ohshima, Y., Schafer, J. C., Li, C., Yoder, B. K., Leroux, M. R. and Scholey, J. M. (2007). Sensory ciliogenesis in *Caenorhabditis elegans*: assignment of IFT components into distinct modules based on transport and phenotypic profiles. Mol Biol Cell 18, 1554–69.

64. Oury, J., Liu, Y., Topf, A., Todorovic, S., Hoedt, E., Preethish-Kumar, V., Neubert, T. A., Lin, W., Lochmuller, H. and Burden, S. J. (2019). MACF1 links Rapsyn to microtubule- and actin-binding proteins to maintain neuromuscular synapses. J Cell Biol 218, 1686–1705.

65. Petry, S. and Vale, R. D. (2015). Microtubule nucleation at the centrosome and beyond. Nat Cell Biol 17, 1089–93.

66. Potter, C., Razafsky, D., Wozniak, D., Casey, M., Penrose, S., Ge, X., Mahjoub, M. R. and Hodzic, D. (2018). The KASH-containing isoform of Nesprin1 giant associates with ciliary rootlets of ependymal cells. Neurobiol Dis 115, 82–91.

67. Potter, C., Zhu, W., Razafsky, D., Ruzycki, P., Kolesnikov, A. V., Doggett, T., Kefalov, V. J., Betleja, E., Mahjoub, M. R. and Hodzic, D. (2017). Multiple Isoforms of Nesprin1 Are Integral Components of Ciliary Rootlets. Curr Biol 27, 2014–2022 e6.

68. Radhakrishnan, P., Shukla, A., Girisha, K. M. and Nayak, S. S. (2019). Biallelic c.1263dupC in DOK7 results in fetal akinesia deformation sequence. Am J Med Genet A.

69. Ravenscroft, G., Sollis, E., Charles, A. K., North, K. N., Baynam, G. and Laing, N. G. (2011). Fetal akinesia: review of the genetics of the neuromuscular causes. J Med Genet 48, 793–801.

70. Reiter, J. F. and Leroux, M. R. (2017). Genes and molecular pathways underpinning ciliopathies. Nat Rev Mol Cell Biol 18, 533–547.

71. Rosenbaum, J. L. and Witman, G. B. (2002). Intraflagellar transport. Nat Rev Mol Cell Biol 3, 813–25.

72. Ruba, A., Luo, W., Yu, J., Takao, D., Evangelou, A., Higgins, R., Khim, S., Verhey, K. J. and Yang, W. (2019). The Ciliary Lumen Accommodates Passive Diffusion and Vesicle Trafficking in Cytoplasmic-Ciliary Transport. bioRxiv 704213, doi: https://doi.org/10.1101/704213.

73. Schou, K. B., Pedersen, L. B. and Christensen, S. T. (2015). Ins and outs of GPCR signaling in primary cilia. EMBO Rep 16, 1099–113.

74. Sharma, N., Kosan, Z. A., Stallworth, J. E., Berbari, N. F. **and** Yoder, B. K. (2011). Soluble levels of cytosolic tubulin regulate ciliary length control. Mol Biol Cell 22, 806–16.

75. Shida, T., Cueva, J. G., Xu, Z., Goodman, M. B. and Nachury, M. V. (2010). The major alpha-tubulin K40 acetyltransferase alphaTAT1 promotes rapid ciliogenesis and efficient mechanosensation. Proc Natl Acad Sci U S A 107, 21517–22.

76. Silva, M., Morsci, N., Nguyen, K. C. Q., Rizvi, A., Rongo, C., Hall, D. H. and Barr, M. M. (2017). Cell-Specific alpha-Tubulin Isotype Regulates Ciliary Microtubule Ultrastructure, Intraflagellar Transport, and Extracellular Vesicle Biology. Curr Biol 27, 968–980.

77. Sirajuddin, M., Rice, L. M. and Vale, R. D. (2014). Regulation of microtubule motors by tubulin isotypes and post-translational modifications. Nat Cell Biol 16, 335–44.

78. Stephens, A. D., Liu, P. Z., Banigan, E. J., Almassalha, L. M., Backman, V., Adam, S. A., Goldman, R. D. and Marko, J. F. (2018). Chromatin histone modifications and rigidity affect nuclear morphology independent of lamins. Mol Biol Cell 29, 220–233.

79. Takao, D., Wang, L., Boss, A. and Verhey, K. J. (2017). Protein Interaction Analysis Provides a Map of the Spatial and Temporal Organization of the Ciliary Gating Zone. Curr Biol 27, 2296–2306 e3.

80. Tan-Sindhunata, M. B., Mathijssen, I. B., Smit, M., Baas, F., de Vries, J. I., van der Voorn, J. P., Kluijt, I., Hagen, M. A., Blom, E. W., Sistermans, E. et al. (2015). Identification of a Dutch founder mutation in MUSK causing fetal akinesia deformation sequence. Eur J Hum Genet 23, 1151–7.

81. Verhey, K. J. and Gaertig, J. (2007). The tubulin code. Cell Cycle 6, 2152–60.

82. Vogt, J., Harrison, B. J., Spearman, H., Cossins, J., Vermeer, S., ten Cate, L. N., Morgan, N. V., Beeson, D. and Maher, E. R. (2008). Mutation analysis of CHRNA1, CHRNB1, CHRND, and RAPSN genes in multiple pterygium syndrome/fetal akinesia patients. Am J Hum Genet 82, 222–7.

83. Vogt, J., Morgan, N. V., Marton, T., Maxwell, S., Harrison, B. J., Beeson, D. and Maher, E. R. (2009). Germline mutation in DOK7 associated with fetal akinesia deformation sequence. J Med Genet 46, 338–40.

84. Wang, L. and Dynlacht, B. D. (2018). The regulation of cilium assembly and disassembly in development and disease. Development 145.

85. Wilbe, M., Ekvall, S., Eurenius, K., Ericson, K., Casar-Borota, O., Klar, J., Dahl, N., Ameur, A., Anneren, G. and Bondeson, M. L. (2015). MuSK: a new target for lethal fetal akinesia deformation sequence (FADS). J Med Genet 52, 195–202.

86. Winters, L., Van Hoof, E., De Catte, L., Van Den Bogaert, K., de Ravel, T., De Waele, L., Corveleyn, A. and Breckpot, J. (2017). Massive parallel sequencing identifies RAPSN and PDHA1 mutations causing fetal akinesia deformation sequence. Eur J Paediatr Neurol 21, 745–753.

87. Wloga, D., Joachimiak, E., Louka, P. and Gaertig, J. (2017). Posttranslational Modifications of Tubulin and Cilia. Cold Spring Harb Perspect Biol 9.

88. Wu, H., Xiong, W. C. and Mei, L. (2010). To build a synapse: signaling pathways in neuromuscular junction assembly. Development 137, 1017–33.

89. Zhang, B., Luo, S., Wang, Q., Suzuki, T., Xiong, W. C. and Mei, L. (2008). LRP4 serves as a coreceptor of agrin. Neuron 60, 285–97.

90. Zhang, W., Coldefy, A. S., Hubbard, S. R. and Burden, S. J. (2011). Agrin binds to the N-terminal region of Lrp4 protein and stimulates association between Lrp4 and the first immunoglobulin-like domain in muscle-specific kinase (MuSK). J Biol Chem 286, 40624–30.

