## Supplemental Methods, Figures and Tables for "Centrosome and ciliary abnormalities in fetal akinesia deformation sequence human fibroblasts"

February 24, 2020

**Supplementary figures**

**Figure S1**: **Mitotic FADS cells exhibit abnormal sloppy microtubule spindles, but spindle stability is not affected upon nocodazole treatment.** (**A**) MRC5 and FADS 1 fibroblasts were cold-treated and mitotic cells stained for β-tubulin (green) and DAPI (blue). (**B**) MRC5 and FADS 1 fibroblasts were treated with high (10 μM, complete loss of polymerisation) or low (170 nM, low polymerisation) concentrations of nocodazole and mitotic cells stained for β-tubulin (green) and DAPI (blue). Representative immunofluorescence images are shown. Arrows mark sites of GOLGI mini stacks. Scale bars, 10 $\mu$m.

**Figure S2**: **FADS affects cilia growth, but not cilia resorption.** (**A**) Representative immunofluorescence images of anti-acetylated α-tubulin (ac-tubulin; magenta) and anti-detyrosinated α-tubulin (detyr-tubulin; magenta) staining in serum starved MRC5 FADS fibroblasts grown on cross-bow-shaped micropattern. Actin (green) was visualised by phalloidin-Alexa Fluor 488 and DNA by DAPI (blue). Scale bars, 10 $\mu$m. (**B**) Cilia resorption after release from 48 h serum starvation was comparable in MRC5 and FADS fibroblasts. PC, primary cilia. (**C**) Laminopathies do not coincide with ciliary defects. Atyp progeria, atypical progeria; EDMD, Emery Dreifuss muscular dystrophy; FPLD, Familial partial lipodystrophy; HGPS, Hutchison Gilford progeria syndrome. (**D**) Representative immunofluorescence images of anti-acetylated α-tubulin (ac-tub; green) and anti-detyrosinated α-tubulin (detyr-tub; green) stainings in growing MRC5 and FADS fibroblasts. Detyrosinated α-tubulin stainings show aggregations in FADS cells. Scale bars, 10 $\mu$m.

**Figure S3**: (**A**) MuSK-depleted MRC5 fibroblasts exhibit shorter PCs. MRC5 cells were transfected with a siRNA against MuSK and cells were serum starved for 48 h, fixed, and stained for acetylated α-tubulin (ac-tubulin; green) and Arl13b (green). DNA (blue) was visualised by DAPI staining. Scale bars, 5 $\mu$m. (**B**) MRC5 cells were transfected with the indicated siRNAs and the protein levels of rapsyn and NUP88 were determined by Western blot analysis and (**C-D**) densitometric quantification using Fiji/ImageJ. Data present the mean ±SD of at least three independent experiments. P-value **<0.01, t-test, two-tailed.

**Figure S4**: **Co-localisation of nucleoporins in MRC5 cells with centrosomes and cilia.** (**A**) MRC5 cells were immunostained with anti-NUP214, anti-NUP62, anti-NUP93, and anti-NUP153 antibodies, respectively, (magenta) and anti-γ-tubulin antibodies (green). (**B**) Sub-confluent MRC5 fibroblasts were serum starved for 48 h, fixed, and stained for NUP214, NUP62, NUP93, and NUP98 (magenta) as well as acetylated α-tubulin (ac-tub; green). (**C**) DOK7 (magenta) localises to the axoneme of the PC in sub-confluent MRC5, FADS 1, and FADS 2 fibroblasts that were serum starved for 48 h. Cilia were visualised by anti-acetylated α-tubulin staining (ac-tub; green). (**D**) No cilia association was seen for MuSK (magenta). Cilia were visualised by anti-acetylated α-tubulin staining (ac-tub, green). DNA was visualised by DAPI (blue). Shown are representative confocal images. Scale bars, 10 $\mu$m.

**Figure S5**: (**A**) Proximity ligation assay (PLA) control experiments. MRC5 cells were processed for PLA using either the probes only or the respective antibodies only in order to rule out false positive PLA foci (red). Scale bars, 10 $\mu$m. (**B**) Intercept regression model of Figure 2A. Growth curve analysis was done using multilevel regression technique using R. The intercept model presented here suggests constant differences in proliferation randomly assigned to the different cell lines and the linear model (Fig. 2A) suggests effects of the different cell lines on proliferation. Statistical comparison of these two models revealed that the linear model is our regression model of choice. Data points represent the mean at the specific time point and the point range show the SEM.

Figure S1

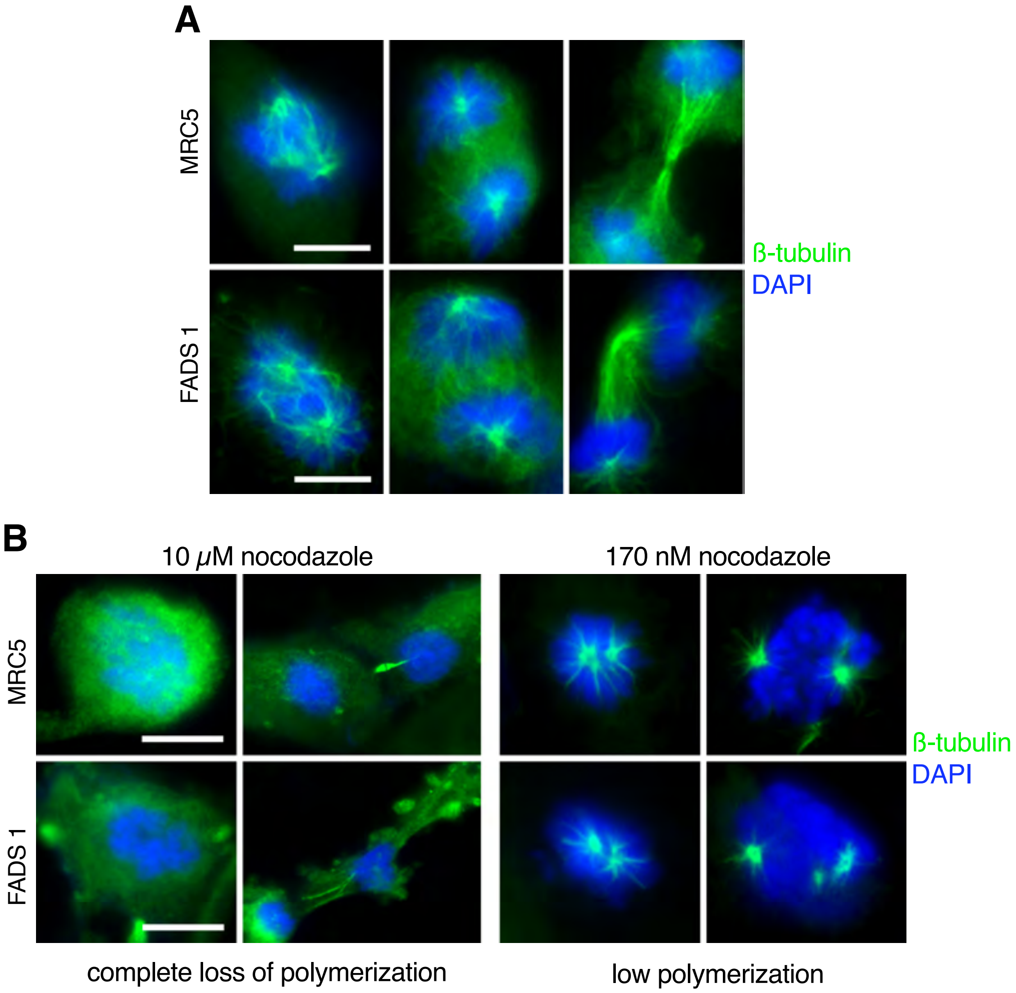
A
MRC5
FADS 1
ß-tubulin
DAPI
B
10 μM nocodazole
170 nM nocodazole
MRC5
FADS 1
ß-tubulin
DAPI
complete loss of polymerization
low polymerization

Figure S2

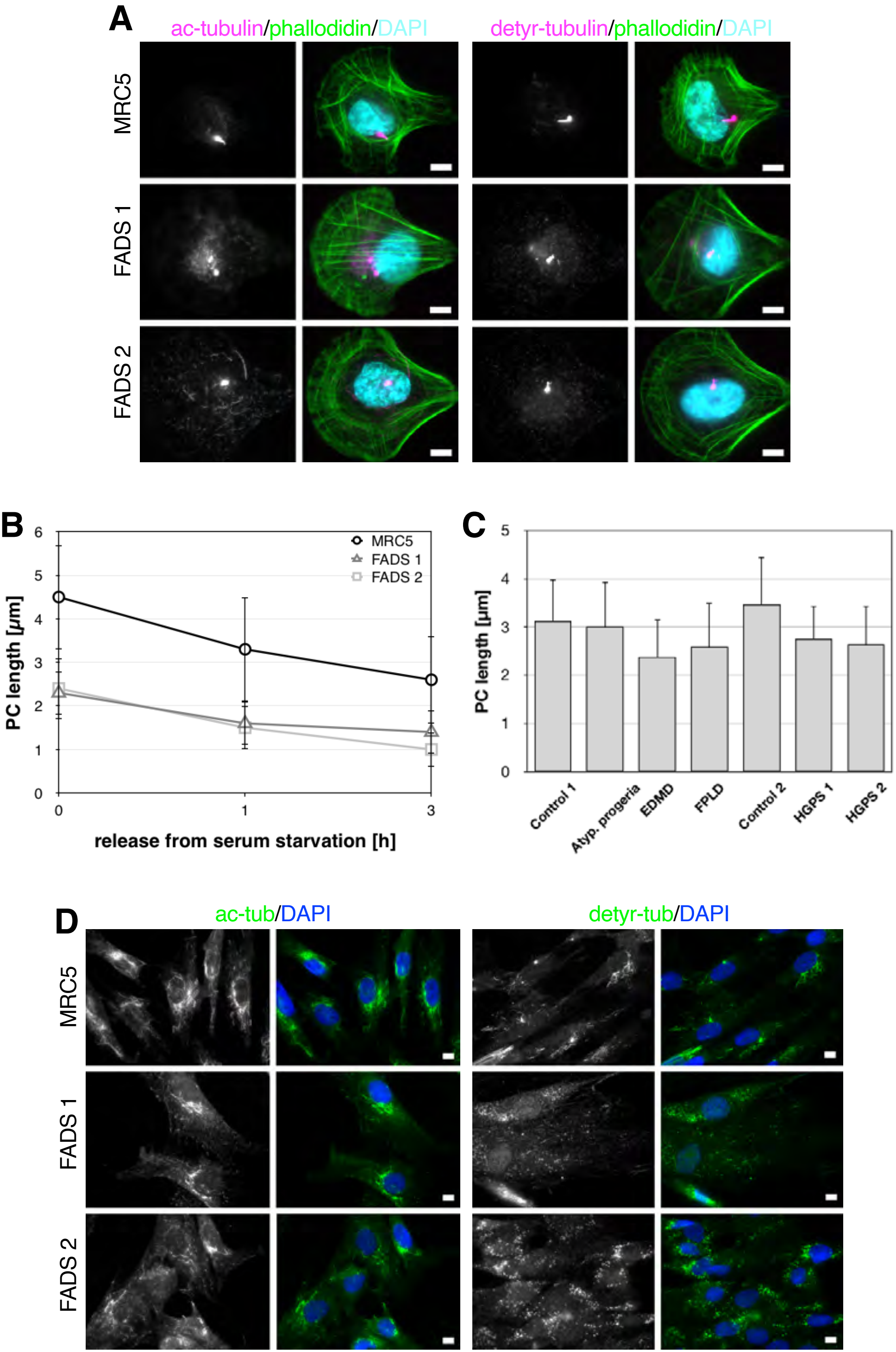
A
ac-tubulin/phallodidin/DAPI
detyr-tubulin/phallodidin/DAPI
MRC5
FADS 1
FADS 2
B
PC length [µm]
0
1
2
3
4
5
6
MRC5
FADS 1
FADS 2
0
1
3
release from serum starvation [h]
C
PC length [µm]
0
1
2
3
4
5
Control 1
Atyp. progeria
EDMD
FPLD
Control 2
HGPS 1
HGPS 2
D
ac-tub/DAPI
detyr-tub/DAPI
MRC5
FADS 1
FADS 2

Figure S3

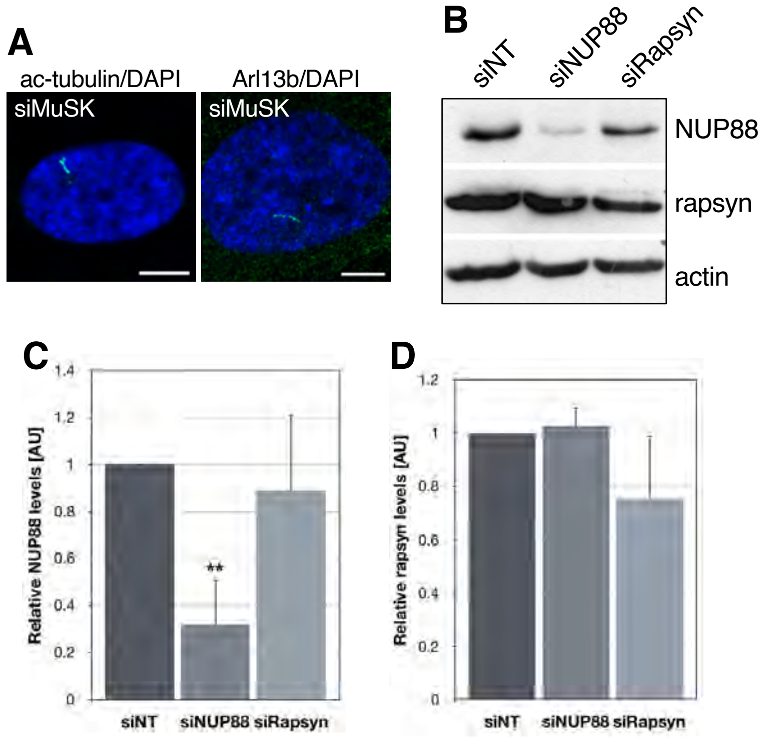
A
ac-tubulin/DAPI
Arl13b/DAPI
siMuSK
siMuSK
B
siNT
siNUP88
siRapsyn
NUP88
rapsyn
actin
C
Relative NUP88 levels [AU]
1.4
1.2
1
0.8
0.6
0.4
0.2
0
**
siNT
siNUP88
siRapsyn
D
Relative rapsyn levels [AU]
1.2
1
0.8
0.6
0.4
0.2
0
siNT
siNUP88
siRapsyn

Figure S4

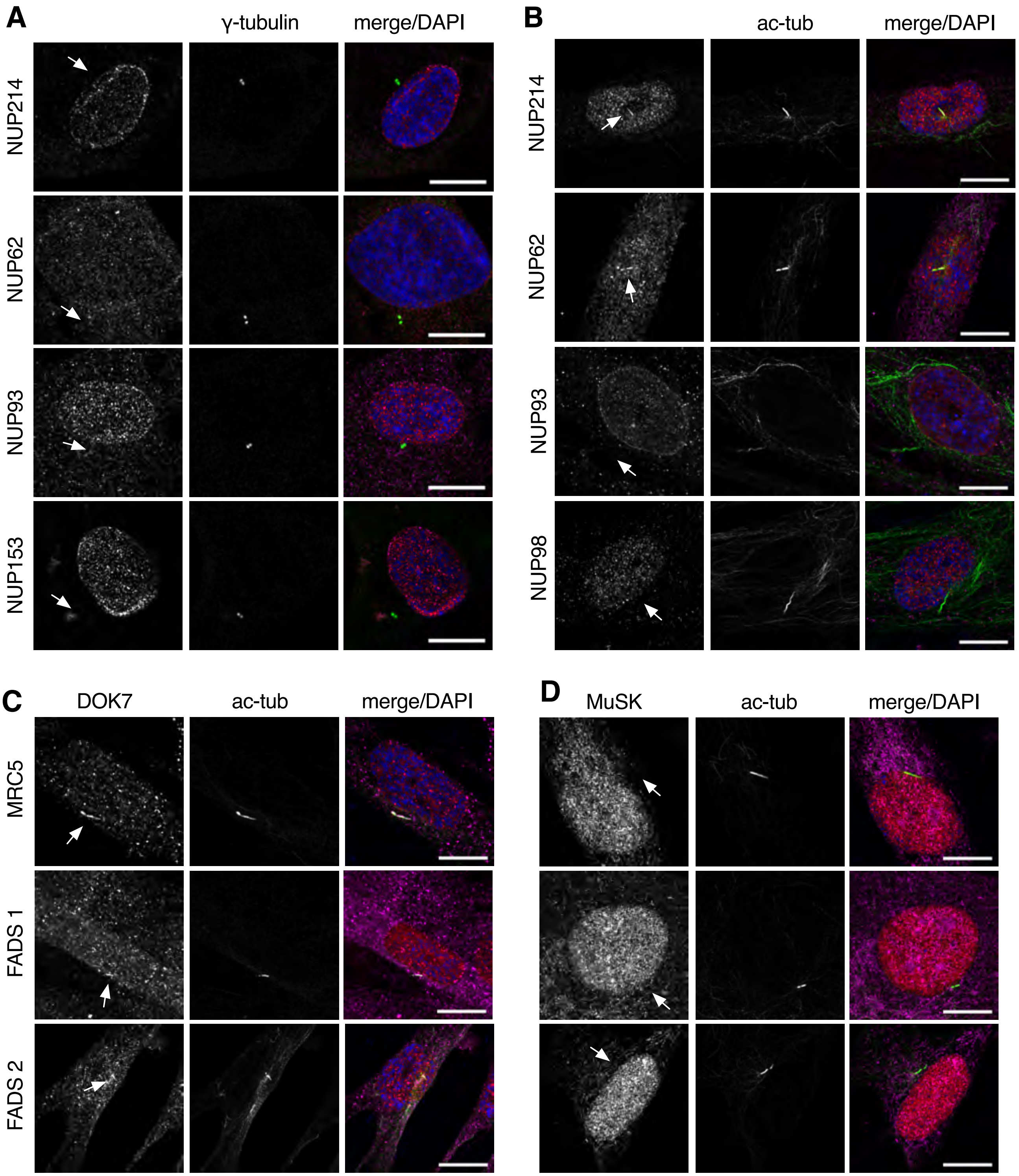
A
γ-tubulin
merge/DAPI
NUP214
NUP62
NUP93
NUP153
B
ac-tub
merge/DAPI
NUP214
NUP62
NUP93
NUP98
C
DOK7
ac-tub
merge/DAPI
MRC5
FADS 1
FADS 2
D
MuSK
ac-tub
merge/DAPI

Figure S5

**A**

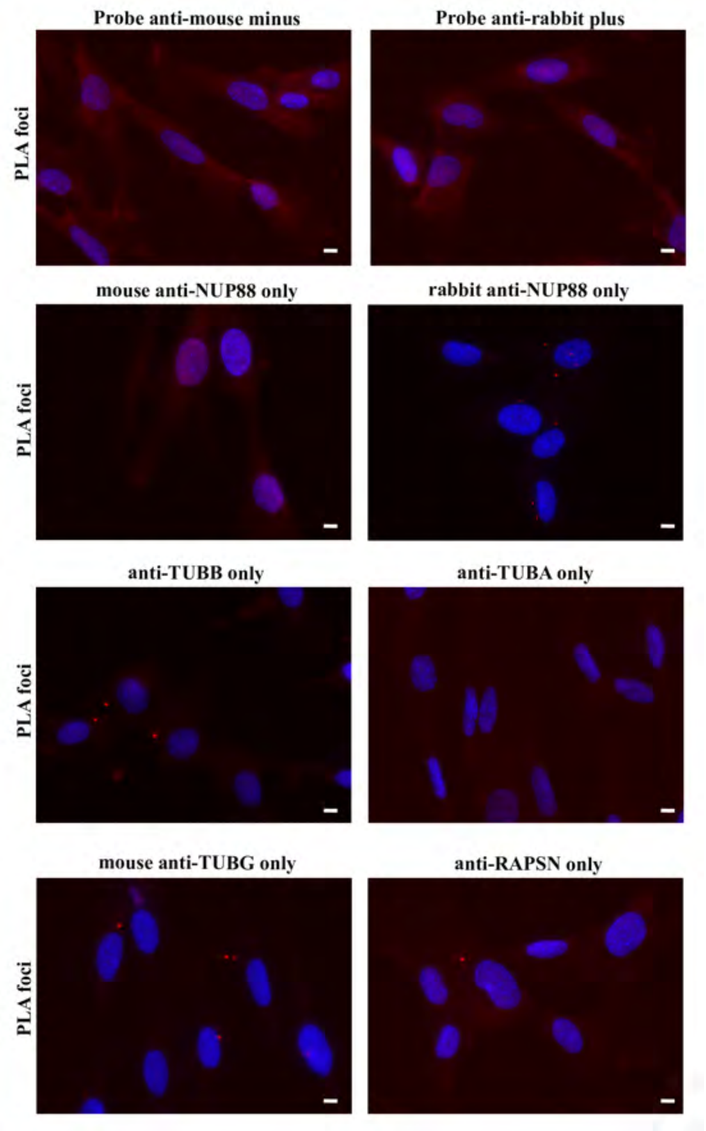

Probe anti-mouse minus
Probe anti-rabbit plus
mouse anti-NUP88 only
rabbit anti-NUP88 only
anti-TUBB only
anti-TUBA only
mouse anti-TUBG only
anti-RAPSN only
PLA foci


**B**

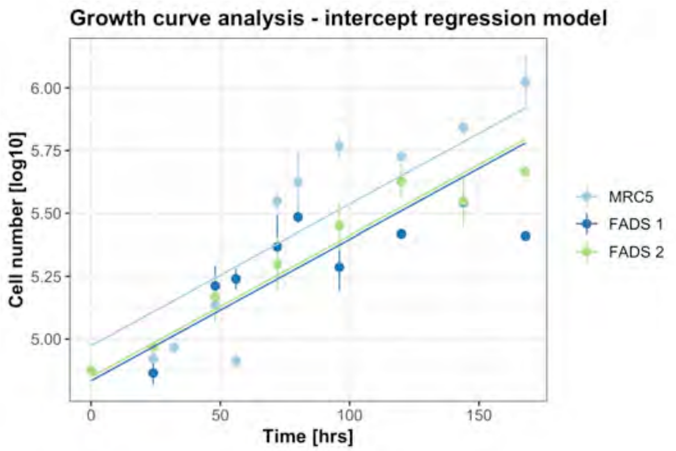

Growth curve analysis - intercept regression model
Cell number [log10]
Time [hrs]
6.00
5.75
5.50
5.25
5.00
0
50
100
150
MRC5
FADS 1
FADS 2

**Table S1:** Cilia localisation of rapsyn and NUP88 in control and FADS fibroblasts.

| | **MRC5** | **FADS 1** | **FADS 2** |
|---|---|---|---|
| rapsyn | 97.4 ± 1.2 | 94.8 ± 2.6 | 97.0 ± 0.9 |
| NUP88 | 94.4 ± 2.9 | 86.9 ± 10.6 | 95.2 ± 1.1 |

Numbers are in percent ± standard deviation. Absolute number of analysed cilia: rapsyn: MRC5, 233; FADS 1, 219; FADS 2, 243; NUP88: MRC5, 200; FADS 1, 289; FADS 2, 146
